# Algal Betaine Triggers Bacterial Hydrogen Peroxide (H_2_O_2_) Production that Promotes Algal Demise

**DOI:** 10.1101/2025.05.07.652642

**Authors:** Delia A. Narváez-Barragán, Lilach Yuda, Dayana Yahalomi, Valeria Lipsman, Sergey Malitsky, Einat Segev

## Abstract

Hydrogen peroxide (H_2_O_2_) plays various roles in the ocean, acting as a signaling molecule at low concentrations and causing oxidative stress when accumulated. While many marine microbes produce H_2_O_2_, its role in microbial interactions remains unclear. Here, we used transcriptomics, genetics, and metabolomics to study H_2_O_2_ dynamics in the interaction between *Emiliania huxleyi* algae and *Phaeobacter inhibens* bacteria. We found that H_2_O_2_ levels rise during algal death and that bacterial H_2_O_2_ production triggers this demise. Manipulating H_2_O_2_ levels shifted the outcome of the interaction. We also uncovered a link between H_2_O_2_ and betaine metabolism: aging algae release betaine, which promotes bacterial H_2_O_2_ production and, in turn, accelerates algal death. Genes involved in H_2_O_2_ and betaine metabolism were upregulated in environmental samples from an algal bloom. Together, our findings identify H_2_O_2_ and betaine as key molecules that modulate algal-bacterial interactions, potentially impacting microbial dynamics in marine ecosystems.

## INTRODUCTION

Reactive oxygen species (ROS) perform a wide range of both harmful and beneficial roles across the tree of life^1^. In the marine environment, hydrogen peroxide (H_2_O_2_) stands out as the most persistent ROS^1,2^. Researchers have detected H_2_O_2_ throughout the water column, from sunlit surface waters to the deep ocean^3,4^, highlighting its stability and ecological relevance in marine systems. Among ROS, H_2_O_2_ uniquely diffuses across cell membranes and persists both inside and outside cells due to its relatively low reactivity^1,2^. Biologically, H_2_O_2_ plays diverse roles: at low concentrations, it functions as a signaling molecule, while at elevated levels, it induces oxidative stress and cellular damage^2^. H_2_O_2_ forms through both abiotic and biotic processes. Abiotic production occurs via photooxidation of dissolved organic matter (DOM) and through rainfall events^4,5^. Biologically, organisms generate H_2_O_2_ as a byproduct of aerobic respiration and photosynthesis. Several sources of extracellular H_2_O_2_ production have been identified in marine systems, including phytoplankton and bacteria^6–9^.

While both phytoplankton and bacteria produce H_2_O_2_, they must tightly regulate its extracellular levels to maintain intracellular redox balance, support essential cellular functions such as cell growth, and prevent oxidative damage^6,7,10^. Some phytoplankton species, however, are highly susceptible to H_2_O_2_ and cannot manage harmful extracellular concentrations on their own. Instead, they depend on coexisting microbes, particularly heterotrophic bacteria, that have evolved strategies to degrade H_2_O_2_ effectively^10,11^. These bacterial detoxification capabilities enhance community-level tolerance in environments with elevated H_2_O_2_ and may create ecological dependencies that influence microbial interactions and shape community composition in the ocean^1,11^. Algal blooms serve as hotspots for H_2_O_2_ production^1,7^, where the high density of microbial cells possibly allows H_2_O_2_ released by one organism to directly affect its neighbors^12^. The rich biomass and DOM produced during blooms enhance both abiotic and biotic H_2_O_2_ generation^1,7^. Heterotrophic bacteria associated with these blooms actively contribute to biotic production^9^. For example, gene expression data suggest that the roseobacter *Ruegeria pomeroyi* produces H_2_O_2_ while degrading dimethylsulfoniopropionate (DMSP), a common algal metabolite abundant during blooms^13^. Bacteria also play a key role in degrading H_2_O_2_, helping to regulate its concentration in the environment^10^. Despite its potential significance in shaping bloom dynamics, microbial communities, and interspecies interactions, the role of H_2_O_2_ in these processes remains poorly understood. The microalga *Emiliania huxleyi* (also known as *Gephyrocapsa huxleyi*^14^) forms dense oceanic blooms that can persist for several weeks before collapsing due to nutrient limitation and viral infection^15,16^. Recent studies have also demonstrated that bacteria can terminate laboratory-induced *E. huxleyi* blooms^17–19^. In natural settings, *E. huxleyi* blooms host diverse bacterial communities, often dominated by members of the roseobacter group. One well-studied species from this group, *Phaeobacter inhibens*, frequently co-occurs with *E. huxleyi* at sea and displays a range of algal-bacterial interactions in laboratory cultures ^17–24^. Initially, *P. inhibens* bacteria promote algal growth, but as the interaction progresses, bacteria shift towards antagonism. *P. inhibens* bacteria induce algal cell death through several mechanisms, including the production of roseobacticides^18^, nitric oxide (NO)^19^, and excessive levels of indole acetic acid (IAA)^17^. Although these are distinct pathogenicity pathways, they all lead to oxidative stress and trigger apoptosis-like programmed cell death in algae^17,19,24^. H_2_O_2_ likely plays a role in this process, as supported by the observation that *P. inhibens* bacteria upregulate genes involved in H_2_O_2_ detoxification in response to *E. huxleyi*-derived *p*-coumaric acid, the precursor of roseobacticides^25^.

Here, we show that H_2_O_2_ accumulates throughout the interaction between *E. huxleyi* and *P. inhibens*, with levels rising significantly during bacterial-induced algal death, suggesting a potential role in triggering algal demise. By manipulating H_2_O_2_ concentrations in co-cultures, we directly influenced the outcome of the interaction. We also identified bacterial genes involved in H_2_O_2_ metabolism and used genetic manipulation to demonstrate their functional contribution to algal death. In addition, we uncovered a previously unrecognized link between H_2_O_2_ and betaine metabolism. Betaine, a well-known algal metabolite is produced by aging algae towards the end of the interaction, and stimulates H_2_O_2_ production in bacteria. Modifying betaine levels altered the trajectory of the algal-bacterial relationship, suggesting that bacteria sense betaine as a proxy of the algal status. Our findings suggest that betaine is an age-related cue from algae, promoting bacterial H_2_O_2_ production and ultimately triggering algal demise.

## RESULTS

### H_2_O_2_ levels impact the algal fate during algal-bacterial interactions

To determine the involvement of H_2_O_2_ in the algal-bacterial interaction we first measured the extracellular H_2_O_2_ levels at different algal growth phases, both in algal mono-cultures and in co-cultures with *P. inhibens* bacteria (Fig. 1A). Cultures were sampled at days 0, 4, 7, 10, and 14 of cultivation, and H_2_O_2_ levels were quantified using the fluorescent probe Amplex Red (see materials and methods). Our data show that algae in mono-cultures continuously produce H_2_O_2_, with levels increasing during the exponential growth phase and decreasing during the late stationary phase (Fig. 1B). However, in co-cultures with bacteria, a different H_2_O_2_ profile is seen over time, with H_2_O_2_ levels decreasing during the early stages of interaction (day 4, Fig. 1A-B) but increasing as bacteria induce algal death (day 14, Fig. 1A-B). These differences in H_2_O_2_ profiles suggest that bacteria impact H_2_O_2_ dynamics throughout the interaction with their algal host.

**Fig 1.**
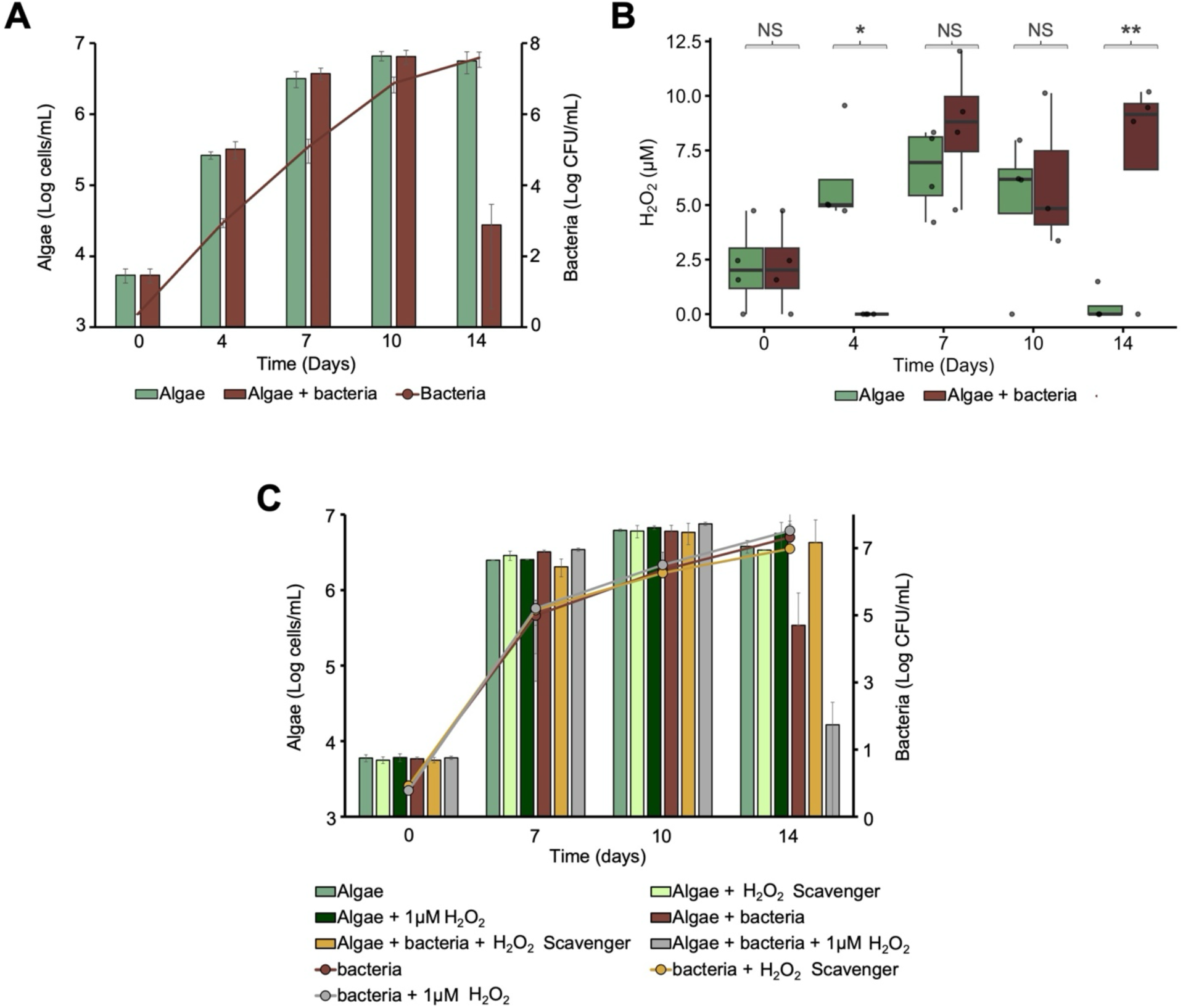
Impacts of H_2_O_2_ levels on algal-bacterial interaction. **A)** *E. huxleyi* algae grown alone (green bars) or in co-cultivation (brown) with *P. inhibens* bacteria (line) along 14 days. **B)** Extracellular H_2_O_2_ concentration (µM) measured at days 0, 4, 7, 10 and 14, in cultures of *E. huxleyi* alone (green) or grown alongside bacteria (brown). **C)** Algae growth alone (green bar) or in co-cultivation (brown bar) with bacteria (lines). Cultures were supplemented with the H_2_O_2_ scavenger Potassium Iodide (1mM KI, light green and yellow) in all time points, or with 1µM H_2_O_2_ (dark green and gray) at days 7, 10 and 12 of the interaction. Error bars represent the standard deviation from the mean of at least 3 biological replicates. Box-plots show results from three independent experiments; black dots indicate individual measurements. Box-plot elements: center line - median; box limits -upper and lower quartiles; whiskers – min and max values. Normality was first assessed using a Shapiro–Wilk test, followed by an unpaired t-test to determine statistical differences between mono- and co-cultures at various time points. NS: Not significant, *p=0.01, **p=0.006.

The involvement of oxidative stress in bacterial pathogenicity towards algae has been documented by us^17,19^ and by others^24,25^. Since H_2_O_2_ can be a prominent inducer of oxidative stress, we investigated whether H_2_O_2_ levels affect the outcome of the algal-bacterial interaction, by manipulating H_2_O_2_ levels. H_2_O_2_ concentrations were either reduced by supplementing cultures with 1 mM potassium iodide (KI), a H_2_O_2_ scavenger^26^, or increased by adding 1 µM H_2_O_2_. Our data show that algae were unaffected by both treatments when grown in mono-cultures (Fig. 1C). However, in co-cultures with bacteria, addition of the scavenger resulted in inhibition of the characteristic algal death, whereas the addition of exogenous H_2_O_2_ accelerated algal death (Fig. 1C). These results demonstrate that the manipulation of H_2_O_2_ levels impacts the algal fate during the algal-bacterial interaction.

### Bacteria modulate the algal H_2_O_2_-related gene expression

The varying levels of H_2_O_2_ during the algal-bacterial interaction could be the result of changes in H_2_O_2_ production, as well as H_2_O_2_ detoxification. Furthermore, these changes could be from either an algal or bacterial source. To understand the algal and bacterial H_2_O_2_ metabolism, we first assessed changes in H_2_O_2_-associated processes in algae through expression analysis of indicative genes. *E. huxleyi* algae are known to produce and metabolize H_2_O_2_^27,28^. To analyze genes involved in algal H_2_O_2_ production and tolerance, we compiled a list of relevant genes by focusing on those associated with enzymatic reactions know to generate H_2_O_2_ either directly or as a byproduct. These genes were identified through literature searches and their annotated functions^7,8,29–32^, as well as homology-based predictions within the MetaCyc database^33^ (table S1). Then, we analyzed the expression patterns of these algal genes in algal mono-cultures and algal-bacterial co-cultures using a RNA-seq dataset that we previously generated^23^. Transcription patterns of algal genes involved in H_2_O_2_ production and degradation were similar between mono and co-cultures during the early and late exponential phases, as well as the stationary growth phase of algal growth (Fig. 2A). During these growth stages, only one gene exhibited different expression patterns, superoxide dismutase (*sod*) was downregulated during the late exponential phase in co-cultures compared to algal mono-cultures (Fig. 2A). However, during algal demise in co-cultures, which corresponds to the late stationary phase in algal mono-cultures, various genes exhibited marked expression differences. In the presence of bacteria, several algal genes that are involved in H_2_O_2_ production were significantly up-regulated, these include *sod* and the genes encoding the reactive oxygen species modulator 1 (*romo1*), spermine oxidase-like (*smo*), FAD-linked sulfhydryl oxidase (*erv_1_*) and NADH dehydrogenases (*ndhs*) from the electron transport chain. The increased expression of these genes suggests increased production of H_2_O_2_, which may be involved in algal demise through the activation of programed cell death^34–37^. Additionally, we observed the downregulation of photosynthesis-related genes, likely due to ROS accumulation^38^. Simultaneously, downregulation of several algal genes involved in H_2_O_2_ detoxification was observed, including genes encoding the glycolate oxidase (*gox*), which is involved in the activation of response genes^39^, protoporphyrinogen oxidase (*ppo*), cytochrome c peroxidase (*cpp*), glutathione peroxidases (*gpx*), and ascorbate peroxidase (*apx*, Fig. 2A). The decreased expression of these genes suggests a compromised algal H_2_O_2_-detoxification process^40,41^. These findings suggest that bacteria enhance the production and decrease the detoxification processes of H_2_O_2_ in algae, thereby potentially accelerating algal demise.

**Fig 2.**
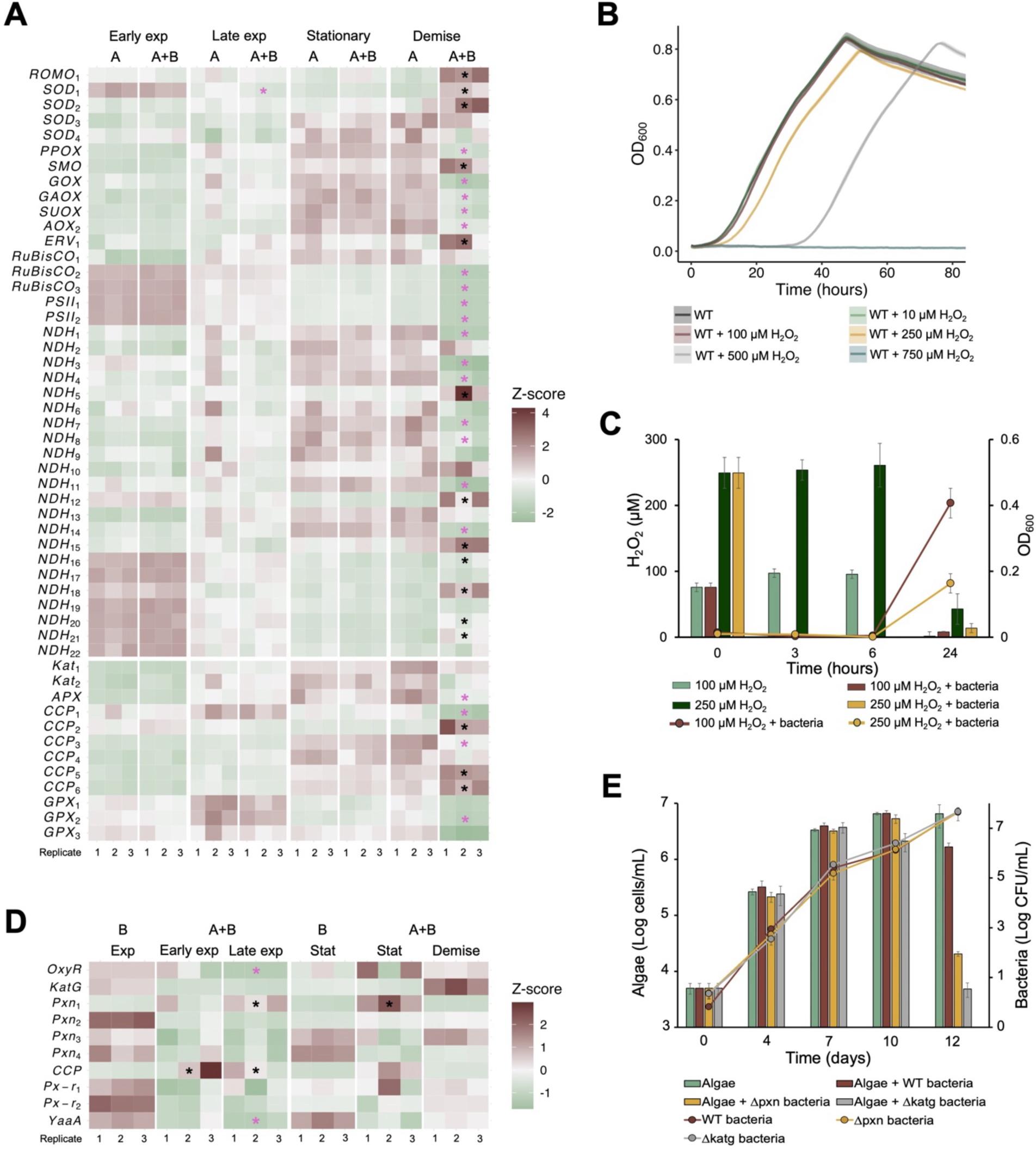
Algal and bacterial H_2_O_2_ production and degradation. **A)** Heatmap showing the expression of algal genes at different growth phases in algal mono-cultures (designated A) or in co-cultures with bacteria (designated A+B). Top panel – H_2_O_2_ production, lower panel - H_2_O_2_ degradation. **B)** *P. inhibens* grown in minimal media supplemented with 0 (black), 10 (green), 100 (brown), 250 (yellow), 500 (gray) and 750 (blue) µM H_2_O_2_. Lines represent the average growth curve based on three biological replicates and shaded areas indicate standard deviation (SD). **C)** Extracellular H_2_O_2_ concentration (µM) measured at 0, 3, 6 and 24 hours of bacterial cultivation, supplemented with 100 (brown) or 250 (yellow) µM H_2_O_2_, compared with media without bacterial addition (light and dark green). Lines represent bacterial growth. **D)** Heatmap showing the expression of bacterial H_2_O_2_ tolerance genes at different growth phases in bacterial pure cultures (designated B) or growing in co-culture with algae (designated B+A). **E)** *E. huxleyi* algae in mono-cultures (green bar) or in co-cultures with *P. inhibens* WT (brown) or *△pxn* (yellow), and *△katg* (gray) mutants. Lines represent bacterial growth. In **A** and **D** heatmaps expression intensity is shown as Z score. Asterisks indicate significant differential gene expression (adjusted *p*-value <0.05), black asterisk indicate upregulation while pink asterisk downregulation. Gene descriptions and log2 fold changes are available at table S1-2. The underlying dataset was previously described in Sperfeld & Narvaez-Barragan *et al*.^23^. Exp: exponential, stat: stationary. In **C** and **E** error bars represent the standard deviation from the mean of 3 biological replicates.

### Bacteria can produce and degrade H_2_O_2_

In addition to the algal impact on H_2_O_2_ levels in co-cultures, bacteria may also contribute to the observed patterns. To test the bacterial influence on H_2_O_2_ levels, we evaluated the ability of *P. inhibens* to metabolize extracellular H_2_O_2_. First, to mimic conditions in which bacteria experience extracellular H_2_O_2_ from an algal source, we examined bacterial growth under increasing concentrations of extracellular H_2_O_2_ in pure bacterial cultures with defined media. Our data show that *P. inhibens* bacteria can grow in H_2_O_2_ concentrations as high as 500 µM (Fig. 2B). Next, to track the bacterial ability to degrade extracellular H_2_O_2_, we cultivated pure bacterial cultures in defined media supplemented with 100 µM or 250 µM H_2_O_2_ and monitored the H_2_O_2_ levels in these cultures. We observed that under both H_2_O_2_ concentrations, the extracellular H_2_O_2_ was completely depleted after 3 hours, as compared to a control sample that did not contain bacteria (Fig. 2C). After 24 hours, bacterial growth was evident, and H_2_O_2_ was detected in bacterial cultures but not in the controls, where H_2_O_2_ was abiotically degraded^1^. These findings suggest that *P. inhibens* bacteria can both degrade and produce H_2_O_2_.

To understand the bacterial mechanisms involved in H_2_O_2_ metabolism in the presence of algae, we analyzed the expression patterns of relevant bacterial genes. First, we compiled a list of genes that encode enzymes with established roles in the bacterial response towards H_2_O_2_^41^ (table S2). Expression analyses of these genes in bacterial pure cultures versus co-cultures with algal partners^23^ revealed several genes encoding peroxiredoxins (*pxn*) that were expressed exclusively in the presence of algae (for example *pxn_1_*, Fig. 2D and table S2). A similar expression pattern was observed for *ccp* (Fig. 2D). Both *pxn* and *ccp* have been reported to be involved in the cell antioxidant defense^42,43^. Additionally, *oxyR*, encoding a H_2_O_2_ sensor^44^ and *katG*, encoding the major catalase involved in H_2_O_2_ degradation^45,46^, were both upregulated in bacteria during the algal demise (Fig. 2D).

To corroborate the involvement of the differentially expressed genes in the bacterial tolerance to H_2_O_2_, we deleted the genes *pxn_1_* and *katG,* and challenged the mutants with different H_2_O_2_ concentrations (fig. S1). Our results show that the *ΔkatG* mutant was the most affected, exhibiting delayed growth under 100mM H_2_O_2_ (fig. S1A) compared to the wild-type (WT), which was not affected by the same H_2_O_2_ concentration (Fig. 2B). In contrast, the *Δpxn_1_* mutant was affected only under resource-limited conditions, such as reduced carbon source concentrations (fig. S2B and E). Limited nutrient bioavailability was previously reported to exacerbate bacterial susceptibility to H_2_O_2_^1,47^. These findings highlight the importance of KatG for H_2_O_2_ detoxification in *P. inhibens* and suggest that bacteria can activate different mechanism to respond to H_2_O_2_ under various environmental conditions, such as variations in resource availability.

Given the compromised ability of the mutant strains to degrade H_2_O_2_, we hypothesized that these mutants would exhibit different dynamics during their interaction with algae. We therefore cultivated mutant bacteria in co-cultures with algae and monitored the bacterial and algal growth (Fig. 2E). Our data show that co-cultures with the mutant strains *Δpxn* and *ΔkatG* exhibit accelerated algal death (Fig. 2E). As both Pxn and KatG are involved in H_2_O_2_ degradation, their absence might result in H_2_O_2_ accumulation, resulting in expedited algal death. To explore this possibility, we measured the H_2_O_2_ levels in co-cultures with the mutant strains, specifically at day 12, which represents the stage of algal demise in co-cultures with mutant bacteria. Our results indicate higher H_2_O_2_ concentrations in the co-cultures with *Δpxn* and *ΔkatG* mutants compared to co-cultures with WT strains (fig. S2). This suggests that impaired bacterial H_2_O_2_ degradation leads to accumulation of H_2_O_2_ in the context of the algal host, which accelerates algal death. These dynamics are similar to the effects observed upon addition of exogenous H_2_O_2_ to algal cultures (Fig. 1C), highlighting the importance of maintaining balanced H_2_O_2_ production and degradation processes during algal-bacterial interactions.

### H_2_O_2_ from a bacterial source induces algal demise

Various marine bacteria possess the genetic potential to produce H_2_O_2_^8,9^, but whether they contribute to the extracellular H_2_O_2_ pool during their interactions with algal partners is unknown. To investigate bacterial H_2_O_2_ production, we monitored the levels of bacterially produced H_2_O_2_ in pure bacterial cultures and in the context of algae. Distinguishing between the bacterial and algal H_2_O_2_ sources in co-cultures is experimentally challenging. Therefore, we cultivated bacteria in 75% algal spent media (see materials and methods), allowing bacteria to sense the chemical repertoire of their algal host, but eliminating the active production of algal H_2_O_2_ during cultivation. Our results revealed that H_2_O_2_ levels increased over time in bacterial cultures that were supplemented with algal spent media but not in control bacterial cultures that were cultivated in a defined medium (Fig. 3A), suggesting that bacteria produce H_2_O_2_ in the context of algae.

**Fig 3.**
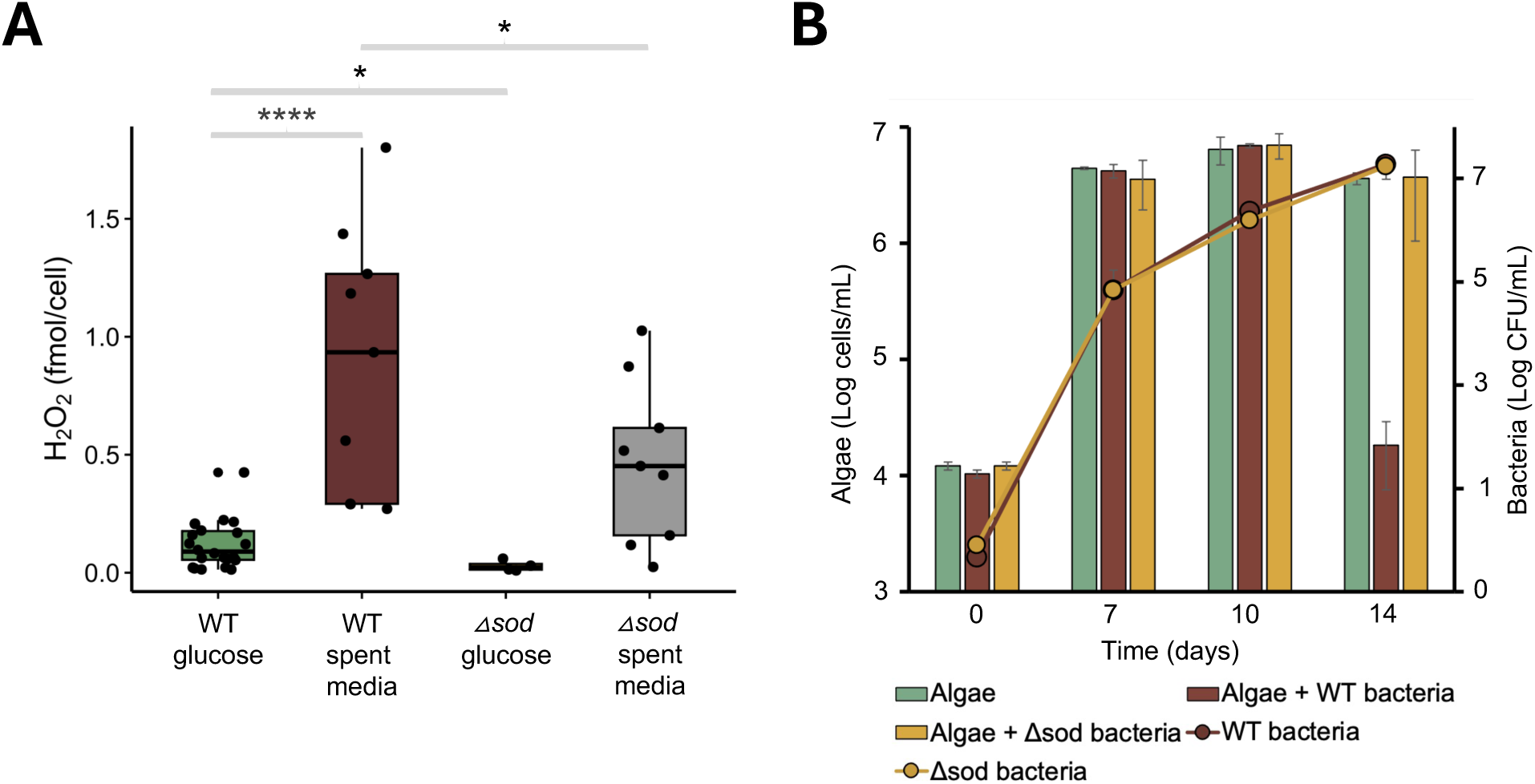
H_2_O_2_ produced by bacteria promotes algal death. **A)** Extracellular H_2_O_2_ concentration (fmol/cell) measured in exponential *P. inhibens* bacteria grown in minimal media with 5.5 mM glucose as a carbon source (WT: green and *Δsod* mutant: yellow), or 75% spent media collected from algae at day 14 (brown). Box-plots show results from three independent experiments; black dots indicate individual measurements. Box-plot elements: center line - median; box limits -upper and lower quartiles; whiskers – min and max values. Normality was first assessed using a Shapiro–Wilk test, followed by a Mann-Whitney test to determine statistical differences between mono- and co-cultures at various time points. *P≤0.03. **B)** *E. huxleyi* grown alone (green bar) or in co-cultivation with *P. inhibens* WT (brown) or *△sod* mutant (yellow). Lines represent bacterial growth. Error bars represent the standard deviation from the mean of at least 3 biological replicates.

To corroborate the bacterial H_2_O_2_ production during co-cultures, we wanted to perturb the bacterial ability to generate H_2_O_2_ and monitor the algal-bacterial interaction. Therefore, we deleted the bacterial gene that encodes a super oxide dismutase (*sod*, table S2) which is essential in the conversion of a superoxide anion to H_2_O_2_ and oxygen (2O_2_^•−^+2H^+^ → H_2_O_2_+O_2_)^48^. The deletion of this gene resulted in a delayed bacterial growth when grown in pure cultures with glucose as a carbon source (fig. S3A), possibly due to the role of superoxide in regulating growth^49^. Interestingly, these effects were less pronounced, or not observed at all, when the *Δsod* mutant was grown in algal spent media (fig. S3B) or in co-cultures with algae (Fig. 3B). Monitoring H_2_O_2_ levels in pure bacterial cultures normalized by cell numbers, showed that the *Δsod* mutant produced less H_2_O_2_ than the WT when cultivated in defined media or algal spent media (Fig. 3A). Furthermore, in co-cultures of the *Δsod* mutant with algae, a delayed algal death was observed (Fig. 3B), suggesting that bacterially produced H_2_O_2_ contributes to promoting algal demise.

### Betaine induces bacterial H_2_O_2_ production

To understand the process of bacterial H_2_O_2_ production during the algal-bacterial interaction, we aimed to identify the elements that regulate H_2_O_2_ production in *P. inhibens*. It was previously suggested that H_2_O_2_ might be produced by the degradation of DMSP. This was based on the co-expression of *dmdA,* the gene encoding the primary DMSP demethylase, and *katG* in environmental samples^13^. However, in our transcriptomic data^23^, we did not observe a correlation between the expression of *katG* and the genes encoding the DMSP-degrading enzymes DmdA^50^ or Bmt^55^ (table S3). Therefore, we explored other genetic modules that are potentially involved in bacterial H_2_O_2_ production (table S2). We identified three operons encoding sarcosine oxidases (*sox*_1_, *sox*_2_ and *sox*_3_, Fig. 4A) which, according to previous reports, can release H_2_O_2_ as a byproduct during the degradation of betaine^51,52^. Betaine is an algal-secreted metabolite which is abundant in marine environments^53^. While the enzymatic machinery responsible for betaine degradation in *P. inhibens* has not been fully elucidated^54^, we observed that these bacteria can utilize betaine as a sole carbon source, although reaching lower yields compared to growth with glucose (fig. S4). Furthermore, *P. inhibens* bacteria exhibit various physiological responses to betaine^22,23,55^, suggesting that they possess the capacity to metabolize this compound. Interestingly, examining the genomic context of the *sox* operons revealed additional nearby elements related to H_2_O_2_ production, such as the *sod* gene, which is a source of H_2_O_2_ in *P. inhibens* (Fig. 3), lactate dehydrogenase (*ldh*) that can form H_2_O_2_^56^, as well as additional components involved in betaine degradation (fig. S5). This genomic proximity suggests a possible link between H_2_O_2_ production and betaine metabolism.

**Fig 4.**
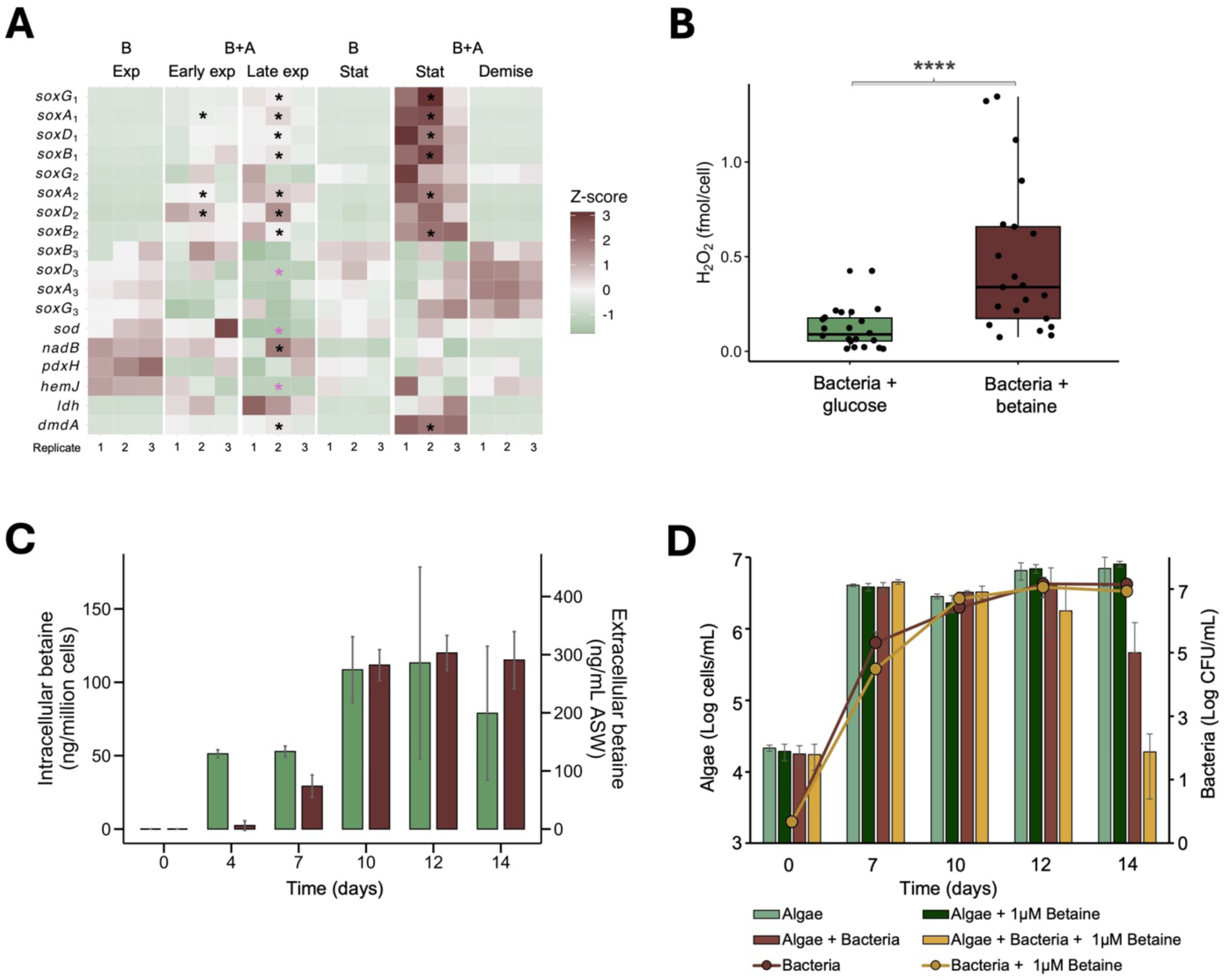
Betaine induces bacterial H_2_O_2_ production driving algal demise. **A)** Heatmap showing the expression of bacterial genes involved in H_2_O_2_ production at different growth phases in bacterial pure cultures (designated B) or in co-culture with algae (designated B+A). Expression intensity is shown as Z score. Asterisks indicate significant differential gene expression (adjusted *p*-value <0.05), black asterisk indicate upregulation while pink asterisk downregulation. Gene descriptions and log2 fold changes are available at table S2. The underlying dataset was previously described in Sperfeld & Narvaez-Barragan *et al*.^23^. Exp: exponential, stat: stationary. **B)** Extracellular H_2_O_2_ concentration (fmol/cell) measured in pure bacterial cultures of WT bacteria cultivated in minimal media using glucose (green) or betaine (brown) as carbon source. Box-plot elements: center line - median; box limits -upper and lower quartiles; whiskers – min and max values. Normality was first assessed using a Shapiro–Wilk test, followed by a Mann-Whitney test to determine statistical differences between mono- and co-cultures at various time points. *P=0.0001. **C)** Intra-(green bars) and extra-cellular (brown bars) betaine levels produced by algae in monocultures, measured on days 0, 4, 7, 10, 12 and 14 of cultivation using liquid chromatography–mass spectrometry (LC–MS). **D)** Algae in monocultures (green bars) or in co-cultures with bacteria (brown bars) supplemented at days 7, 10 and 12 with 1 μM betaine (dark green bars for monocultures and gray bars for co-cultures). Lines represent bacterial growth. In **C** and **D** error bars represent the standard deviation from the mean of at least 3 biological replicates.

Therefore, we examined the possible involvement of betaine in bacterial H_2_O_2_ production. First, we examined the expression levels of the *sox* operons in pure bacterial cultures and in algal-bacterial cultures. Our data revealed that two of these operons, *sox*_1_ and *sox*_2_, are highly expressed during interaction with algae but not when bacteria are grown in pure cultures (Fig. 4A and table S2). To explore whether specifically betaine influences H_2_O_2_ production in bacteria, we monitored H_2_O_2_ levels in pure bacterial cultures cultivated in defined media with either glucose or betaine as a sole carbon source. We maintained the same carbon concentration across experiments and normalized the results by bacterial cell numbers. Our results indicate a roughly 2-fold increase in the levels of H_2_O_2_ in cultures that were cultivated with betaine (Fig 4B). Next, we analyzed the impact of betaine on the relative expression of genes that are putatively involved in bacterial H_2_O_2_ production. Therefore, pure bacterial cultures were cultivated with betaine or glucose as a sole carbon source, RNA was extracted, and expression levels were compared between cultures via qRT-PCR. According to our results, betaine strongly induces *sox*_1_ expression (24-log_2_ fold) but not *sox*_2_ (table S4), suggesting that *sox*_1_ may play a key role in betaine and H_2_O_2_ metabolism in *P. inhibens.* However, as mentioned before, both *sox* operons were upregulated in the presence of algae (Fig. 2A). Additional H_2_O_2_-producing genes were also upregulated in response to betaine; *sod*^48^ (2-log_2_ fold), pyridoxamine 5’-phosphate oxidase (*pdxh*^57^, 0.5-log_2_ fold) and protoporphyrinogen oxidase (*hemJ*^58,59^, 1-log_2_ fold) (Fig. S6B). Overall, our data suggest that bacterial H_2_O_2_ production is promoted by extracellular betaine.

As betaine is a common algal metabolite^53^ that appears to impact bacterial H_2_O_2_ metabolism, we wished to characterize betaine levels in algal cultures of *E. huxleyi.* To this end, cells and supernatants were collected from algal mono-cultures on days 0, 4, 7, 10, 12 and 14, and subjected to intra- and extracellular betaine measurements using liquid chromatography–mass spectrometry (LC–MS) analysis (Fig. 4C). Our data demonstrate that algae produce intracellular betaine throughout algal growth and secrete increasing levels of betaine as the algal culture ages, during the stationary phase. Interestingly, in algal-bacterial co-cultures, algal demise due to bacteria is seen at day 14 of incubation (Fig. 1A) and appears to be preceded by a period in which algal betaine secretion is constantly high, as evident in algal mono-cultures (Fig. 4C).

Next, we evaluated whether the concentration of secreted betaine observed in algal mono-cultures can drive bacterial pathogenicity, likely through bacterial H_2_O_2_ production. Therefore, we calculated based on our LC-MS measurements that the average concentration of extracellular betaine in the stationary phase is 2.5 μM. Consequently, we supplemented algal-bacterial co-cultures with 1μM betaine at days 7, 10 and 12, time points at which algal demise is not yet seen in co-cultures. Betaine supplementation triggered earlier algal death compared to non-supplemented controls (Fig. 4D). In contrast, axenic algal cultures supplemented with betaine did not show signs of cell death (Fig. 4D), indicating that the effect depends on the presence of bacteria. These results suggest that betaine secretion by aging algae triggers bacterial H_2_O_2_ production, which in turn can contribute to algal demise.

### Co-expression of bacterial H_2_O_2_ and betaine related genes in the environment

The novel metabolic connection between betaine and H_2_O_2_ uncovered in this study could have environmental significance. Given that *E. huxleyi* algae and *P. inhibens* bacteria naturally co-occur in oceanic ecosystems^17,20^, and considering that other algal-associated bacteria possess similar genomic capabilities,^9,60,61^ our findings may hold ecological relevance. However, bridging between simplified laboratory cultures and the complex marine environment is challenging. Yet, once a mechanism is revealed in a model system, indicative proxies can be measured at sea. Therefore, as a first step in elucidating the possible relevance of betaine-induced H_2_O_2_ metabolism at sea, we explored the transcript abundance of bacterial H_2_O_2_ and betaine-related genes in algal-rich ocean environments. Analysis of a metatranscriptomic dataset from the TARA Oceans expedition^62^ showed a positive correlation between chlorophyll *a* (as an indicator of algal abundance), transcript abundance of bacterial genes related to betaine degradation (*mttb* and *soxD,G*), and transcript abundance of bacterial genes related to H_2_O_2_ production (*hemJ* and *ldh*) (fig. S6). These observations suggest that bacterial betaine degradation and H_2_O_2_ production are possibly enhanced in algal-rich environments.

Next, we analyzed the expression patterns of bacterial genes related to H_2_O_2_ degradation, H_2_O_2_ production, and betaine metabolism during a natural algal bloom. Algal blooms naturally occur in the ocean due to an influx of inorganic nutrients and low abundance of predators reaching algal densities of at least 10^6^ cells/L^63^. We re-analyzed a previously published transcriptomics dataset that followed bacterial gene expression during an algal bloom^64^. Our analysis revealed increased expression of genes involved in H_2_O_2_-degradation genes during the second half of the bloom (April 6th and 9th timepoints), with some genes maintaining high expression until algal demise (Fig. 5). These data suggest that bacteria experience elevated H_2_O_2_ levels during later phases of the algal bloom. Interestingly, the higher expression of bacterial H_2_O_2_-related genes appears to be independent of algal abundance. Altought algal growth is evident by March 26th and April 3rd, the induction of bacterial H_2_O_2_-degrading genes is not observed until April 6th. In line with this observation, bacterial H_2_O_2_-producing genes follow a similar trend, showing increased expression toward the end of the bloom, with *pdxH*, *hemJ* and *sod* exhibiting increased expression levels (Fig. 5). This enhanced expression suggests a bacterial contribution to H_2_O_2_ levels during the algal bloom. Notably, in our experimental system, these H_2_O_2_-producing genes were also induced by betaine (table S6), supporting a mechanistic link. Expression of genes in the sox operon, responsible for betaine-dependent H_2_O_2_ production, was observed before and during algal demise, with *sox*G showing enhanced expression (Fig. 5). Genes involved in bacterial betaine degradation were expressed throughout the bloom, possibly reflecting the dual role of betaine as carbon an nitrogen source^61^. This may explain the expression of betaine-related genes at the beginning of the bloom, consistent with observations that *P. inhibens* and other marine bacteria metabolize betaine to resume growth^23^. High expression of bacterial betaine-related genes was also observed at the end of the bloom. However, it is important to note that the sampled bloom was biphasic; an influx of nutrient-rich coastal waters on April 20th led to a second bloom phase^64^. The overlap between the demise of the first bloom and the onset of the second bloom may complicate interpretation of bacterial gene expression patterns at this timepoints.

**Fig 5.**
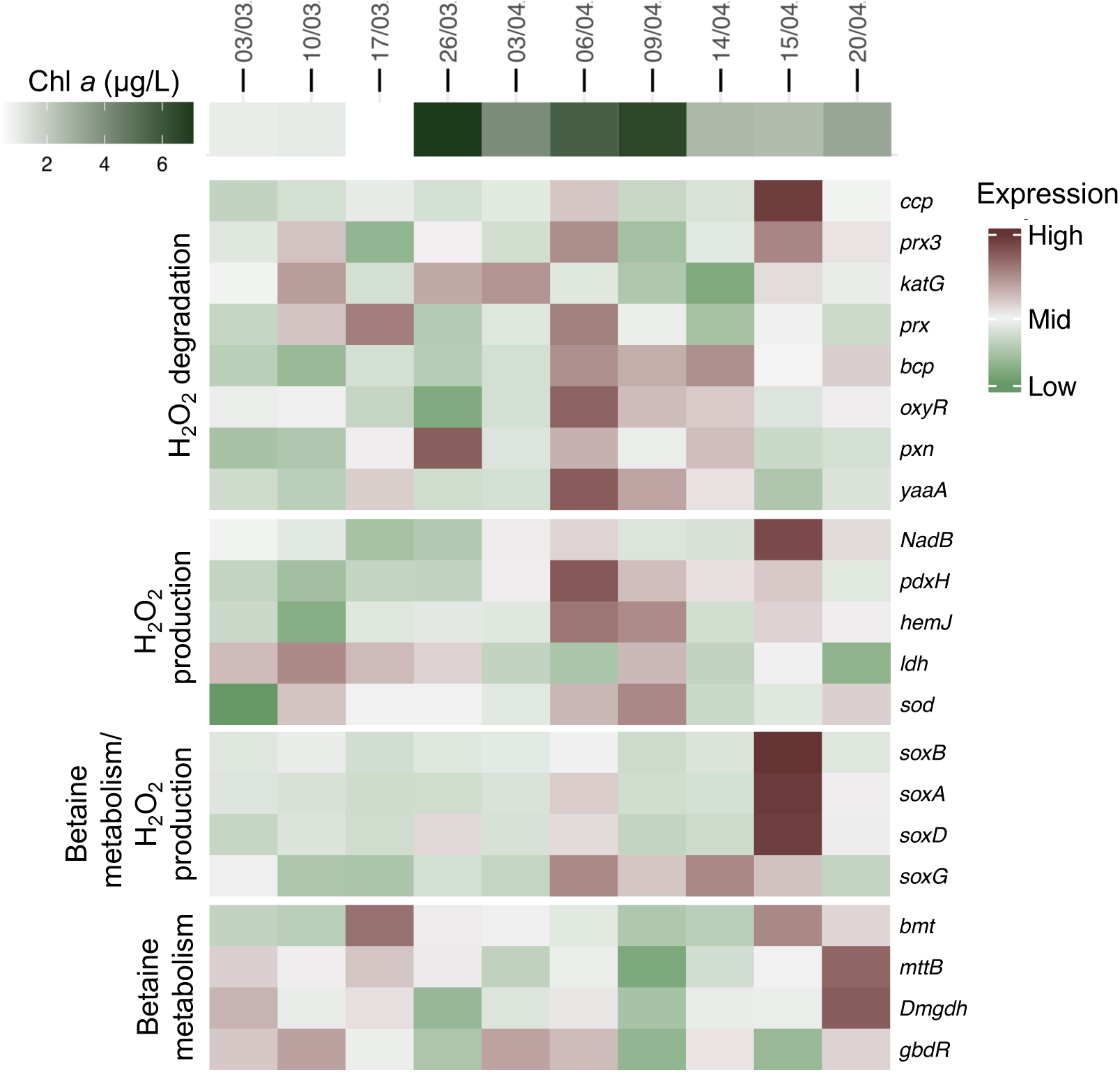
Co-expression of bacterial H_2_O_2_ and betaine-related genes during a natural algal bloom. Expression profiles of 21 bacterial genes involved in H_2_O_2_ degradation, H_2_O_2_ production, and betaine metabolism across 10 time points during an algal bloom event (March to April 2020). Transcript abundance is presented as a Z-score normalized values. Algal abundance, represented by chlorophyll *a* (Chl *a*) concentration, is shown in the green heatmap in the top of the figure alongside the sampling days. Data are based on metatranscriptomic analysis from Sidhu *et al*.^64^.

Interestingly, the first bacterial enzymes in the pathway of betaine degradation, *mttb*^65^ and *bmt*^55^, along with the transcriptional activator *gbdr*^66^, were expressed early in the bloom (March 26th and April 3rd timepoints), followed by the induction of *sox* genes, and then by increased expression of genes involved in H_2_O_2_-production during the bloom decline. This pattern suggests that bacterial betaine metabolism may precede and contribute to oxidative stress, ultimately playing a role in bloom collapse.

Together, these environmental observations complement our laboratory findings, suggesting that bacterial betaine degradation and the associated H_2_O_2_ production may influence the algal fate during natural bloom events, highligthing the broader ecological relevance of this metabolic connection.

## DISCUSSION

### Modulating H_2_O_2_ levels changes the fate of algal-bacterial interactions

Interactions between algae and bacteria rely on the exchange of nutrients and metabolites^67^. However, inorganic molecules are often overlooked in the context of this microbial communication, despite their ability to influence and shape microbial interactions^19^. Among these inorganic compounds, H_2_O_2_ appears to markedly impact the *E. huxleyi-P. inhibens* interaction. H_2_O_2_ levels measured during the interaction change in response to bacterial presence (Fig 1A-B), and manipulating H_2_O_2_ levels, either by decreasing or increasing them, determines whether algae and bacteria will coexist or whether the algal population will collapse, respectively (Fig. 1C).

Bacteria have the potential to act as key regulators of H_2_O_2_ levels in natural aquatic systems^9^. We found that *P. inhibens* bacteria possess multiple strategies to detoxify H_2_O_2_ (Fig.2B-D). Disrupting the bacterial H_2_O_2_ degradation capacity has direct consequences for the interaction, particularly impacting the fate of the algal partner (Fig. 2E). Consistent with environmental observations in other heterothropic bacteria^46^, KatG appers to function as the primary enzyme for H_2_O_2_ degradation in *P. inhibens* (fig. S1). Although the *Δkatg* mutant shows impaired growth in minimal media supplemented with H_2_O_2_, its growth in co-culture with algae is comparable to the WT (Fig. 2E), suggesting that other detoxification mechanisms may compensate to maintain low intracelular bacterial H_2_O_2_ concentrations^10^. We detected peroxiredoxin (*pxn1*) expression only in the presence of the algal host (Fig. 2D), indicating that H_2_O_2_-degrading enzymes may exhibit functional redundancy that is dependent on the environmental context. Aditionally, *pxn1* is further influenced by nutrient limitation coditions (fig. S1). This ability to maintain balanced redox in response to multiple environmental challenges, also observed in *Vibrio vulnificus* and *E. coli*^68,69^, may provide a competitive advantage for marine bacteria constantly exposed to fluctuating H_2_O_2_ levels^4^.

In contrast, algal H_2_O_2_-degrading enzymes expression does not appear to change in response to bacterial presence during exponential growth and the early stationary phase. However, during algal demise, different enzymes involved in H_2_O_2_ degradation, including *apx*, *ccp1,3*, and *gpx 2-3* are downregulated (Fig. 2A). This response suggests that an impaired algal H_2_O_2_-detoxification process might be part of the bacterial pathogenic strategy^40^, contributing to H_2_O_2_ accumulation and redox imbalance that leads to algal death. Additionally, H_2_O_2_ homeostasis may become incleasingly important as the algal host ages. Indeed, in the bloom-forming diatom *Coscinodiscus radiatus,* ROS scavenging via extracellular vesicles is essential in late growth stages for preventing algal death^15^. These findings underscore the central role of H_2_O_2_ levels in shaping algal-bacterial interactions, showing that redox balance is essential not only for individual fitness but also for determining the outcome of more complex interactions. As human activity continues to elevate oxidative stress in marine environments^1,2^, and H_2_O_2_ is increasingly used to control harmful algal blooms^70^, it becomes impportant to understand how microbes respond to H_2_O_2_ fluctuations.

### Both algae and bacteria contribute H_2_O_2_ during their interaction

Our results suggest a dynamic interplay between H_2_O_2_ production and removal during the *E. huxleyi – P. inhibens* interaction. We observed two distinct shifts in extracellular H_2_O_2_ levels during the algal-bacterial interaction; an initial decrease followed by a later accumulation (Fig. 1B). Algae produce higher levels of H_2_O_2_ during exponential growth, but this production declines as they enter the stationary phase (Fig. 1A–B). During this transition, the expression of H_2_O_2_-degrading enzymes increases (Fig. 2A), while photosynthetic gene expression decreases (Fig. 2A), both indicators of algal aging^15,71^. These results suggest that *E. huxleyi* reduces H_2_O_2_ production as it ages. However, in the presence of bacteria, algae may continue contributing to extracellular H_2_O_2_ pool during their demise, as indicated by the overexpression of algal genes associated with H_2_O_2_ production (Fig 2A).

We also found that *P. inhibens* produces H_2_O_2_, with increasing levels in the presence of the algal host or host-derived metabolites (Fig. 1B and 3A). Interestingly, H₂O₂ appears to function as an important signal for bacterial growth. A Δ*sod* mutant, which produces lower levels of H_2_O_2_, grows more slowly than the WT in defined media (fig. S3A). This growth impairment may result from the accumulation of toxic superoxide (O₂⁻), as previously reported in *sod* mutants of *E. coli* and *Salmonella typhimurium*^72,73^. However, when the *P. inhibens* Δ*sod* mutant was grown in algal spent media containing host-derived H_2_O_2_, its exponential growth was comparable to the WT (fig. S3B). Although, towards the stationary phase, the mutant reached a lower maximal yield, similar to what has been reported for other *sod* mutants, which exhibit increased sensitivity during this phase^72,73^. Likewise, in co-culture with *E. huxleyi*, where H_2_O_2_ is continuously released, the mutant grows at levels comparable to the WT (Fig. 3B). These suggest that external H_2_O_2_ may partially compensate for reduced endogenuos production. Similar responses have been observed in other roseobacters, such as *Ruegeria pomeroyi* and *Roseobacter sp. strain AzwK-3b*., where exposure to O₂⁻ and H_2_O_2_ influence growth and redox regulation^49^. These findings suggest that *P. inhibens,* and potentially other marine bacteria, may constantly secrete H_2_O_2_ as a self-generated signal and increase its production upon encountering a host or host-derived cues.

Our results indicate that both algae and bacteria release H_2_O_2_ into their shared environment and respond to changes in its concentration. This bidirectional exchange suggests a chemical feedback loop in which H_2_O_2_ is not only a metabolic byproduct but also a central component of cross-kingdom inorganic communication.

### H_2_O_2_ from a bacterial source is involved in pathogenicity

In the algal-bacterial interaction studied here, H_2_O_2_ accumulation may be the signal that drives the shift from coexistence to collapse. The Δ*sod* mutant is impaired in H_2_O_2_ production and causes a delayed algal death (Fig. 3), suggesting that bacterial-derived H_2_O_2_ contributes to promoting algal demise. For bacterial H_2_O_2_ to impact neighboring algal cells, close proximity and the formation of gradients within the phycosphere appear to be essential^12,67^. During algal demise, bacterial cells increase their attachment to the algal surface^17^, potentially allowing localized concentrations of inorganic molecules to act as signals that amplify PCD-like responses in algae^16,19,74^. Additionally, some algal genes involved in H₂O₂ production are upregulated primarily in response to bacterial pathogenicity (Fig. 2A). This raises the possibility that bacteria manipulate the algal oxidative metabolism to amplify the H₂O₂ signal and trigger a PCD-like process, as previously observed during *E. huxleyi* viral infections^16^ and upon exposure to the inorganic molecule NO^19^. Interestingly, NO and H₂O₂ are known to act synergistically to modulate responses in plants^75^, and similar interactions may occur in algae, which show comparable sensitivity to oxygen and nitrogen species^19^. Moreover, bacterial roseobacticides that kill the algal host, induce expression of bacterial H₂O₂ defense mechanisms^25^, suggesting that roseobacticides production may be coupled with an increase in H₂O₂ levels. Together, these findings point to the orchestration of multiple bacterial pathways involved in algal death, highlighting the complex molecular interplay underlying bacterial pathogenicity.

Notably, other heterotrophic bacteria that interact with algae also possess the capacity to produce H₂O₂^9^, suggesting that this phenomenon may be widespread in the ocean. As a result, the potential implications of fluctuating H₂O₂ levels, particularly in the context of host-microbe interactions, has likely been underestimated.

### Betaine signals algal aging and triggers a corresponding bacterial response

Bacteria use a variety of algal-derived metabolites, and specific host signals can influence bacterial behavior to promote either mutualistic or pathogenic interactions^17–19,67^. Shifting from mutualism to pathogenicity likely depends on the bacterial ability to sense and respond to cues that reflect the physiological state of the host^19,76^. We found that *E. huxleyi* increases betaine production and secretion as it enters the stationary phase (Fig. 4C), suggesting that betaine levels are correlated with algal aging. In co-culture, *P. inhibens* responded with increased expression of betaine degradation genes during the stationary phase (Fig. 4A). This dose-dependent response suggests that bacteria may sense elevated betaine levels as a cue of algal senescence and adjust their metabolism accordingly. A similar pattern has been observed in the bloom-forming diatom *Coscinodiscus radiatus,* where rising betaine concentrations were associated with the onset of senecense^15^. Interestingly, several algal-derived compounds that are associated with bacterial-induced cell death also accumulate during the late stages of algal growth. These algal compounds include 1) extracellular nitrite, which is reduced by bacteria to NO^19^, 2) *p*-coumaric acid, the degradation product of the algal cell wall and the precursor for roseobacticide biosynthesis^25^, 3) tryptophan, which promotes enhanced bacterial production of indole-3-acetic acid (IAA)^17^, and 4) DMSP which accumulates toward the end of *E. huxleyi* growth and triggers pathogenic responses in specific *Sulfitobacter* strains^76^. When considered alongside these senescence-associated algal metabolites, betaine emerges as a novel link between algal physiology and bacterial pathogenicity. Betaine not only supports bacterial growth (fig. S4), but also influences the expression of bacterial genes involved in promoting algal demise (Fig. 3A and table S4). These findings support that bacteria can sense distinct metabolic fingerprints of their host^77^ to shift their behavior accordingly.

Moreover, we found that bacterial genes involved in H_2_O_2_ and betaine metabolism are co-expressed in association with phytoplankton (fig. S6) and during algal blooms (Fig. 5), suggesting that this form of chemical communication may extend beyond the *P. innhibens* – *E. huxleyi* system. Interestingly, in other microalgae such as *Chrysochromulina* sp. and *Amphidinium carterae,* betaine levels peak earlier, before the stationary phase^78^, suggesting that betaine signaling may vary across taxa.

To conclude, our findings deepen the understanding of the molecular mechanisms that shape algal-bacterial interactions, and highlight H₂O₂ and betaine as key regulatory molecules. This insight is particularly important in the context of climate change, which is expected to alter microbial betaine degradation^53^ and, in turn, impact both betaine and H₂O₂ concentrations in the ocean, potentially influencing microbial dynamics across multiple scales.

## MATERIALS AND METHODS

### Strains and growth conditions

The bacterial strain *Phaeobacter inhibens* DSM 17395 was obtained from the German Collection of Microorganism and Cell Cultures (DSMZ, Braunschweig, Germany). It was grown in either ½ YTSS medium (yeast extract, 2 g/L; trypton, 1.25 g/L; sea salts; 20 g/L), or artificial seawater (ASW) medium based on the protocol of Goyet and Poisson^79^. ASW contained mineral salts (NaCl, 409.41 mM; Na_2_SO_4_, 28.22 mM; KCl, 9.08 mM; KBr, 0.82 mM; NaF, 0.07 mM; Na_2_CO_3_, 0.20 mM; NaHCO_3_, 2 mM; MgCl · 6 H_2_O, 50.66 mM; CaCl_2_, 10.2 mM, SrCl_2_ · 6 H_2_O, 0.09 mM), L1 trace elements (Na_2_EDTA · 2H_2_O, 4.36 mg/L; FeCl_3_ · 6 H_2_O, 3.15 mg/L; MnCl_2_ · 4 H_2_O, 178.1 μg/L; ZnSO_4_ · 7 H_2_O, 23 μg/L; CoCl_2_ · 6 H_2_O, 11.9 μg/L; CuSO_4_ · 5 H_2_O, 2.5 μg/L; Na_2_MoO_4_ · 2 H_2_O, 19.9 μg/L; H_2_SeO_3_, 1.29 μg/L; NiSO_4_ · 6 H_2_O, 2.63 μg/L; Na_3_VO_4_, 1.84 μg/L; K_2_CrO_4_, 1.94 μg/L), L1 nutrients (NaNO_3_, 882 μM; NaH_2_PO_4_ · 2 H_2_O, 36.22 μM), 5 mM NH_4_Cl, 33 mM Na_2_SO_4_, and 5.5 mM Glucose, adjusted to a pH 8.2 with HCl. For especific experiments, the carbon source was replaced with either 1mM glucose or 6.6mM betaine.

*P. inhibens* mutants mutants strains *Δkatg* and *Δpxn* were supplemented with 30 μg/ml gentamycin, and *Δsod* with 150 μg/ml kanamycin. When indicated 1 mM potassium iodide (KI, Sigma-Aldrich) or H_2_O_2_ (Bio-Lab ltd) was added to the ASW medium.

The algal strain *E. huxleyi* CCMP3266 was obtained from the National Center for Marine Algae and Microbiota (Bigelow Laboratory for Ocean Sciences). Algae were cultured in ASW medium at 18 °C under a light–dark cycle of 18 h of light and 6 h of dark, with an illumination intensity of 150 mmol m^−2^ s^−1^. Absence of bacteria in axenic algal cultures was monitored periodically by plating on 1⁄2 YTSS plates and by microscopy.

For experiments using algal spent media, *E. huxleyi* cultures were grown for 14 days under the conditions described above. Cultures were collected and filtered (0.2 µM pore filter,Thermo Fisher Scientific) to remove algal cells. The pH of the resulting spent media was adjusted to 8 and re-filtered to ensure sterility. For bacterial growth experiments, 75% algal spent media was mixed with 25% ASW (v/v, ratio of 3:1), and supplemented with L1 trace elements, L1 nutrients, 5.5 mM glucose, 33 mM Na_2_SO_4_, and 5 mM NH_4_Cl.

### Bacterial growth curves

*P. inhibens* was streaked from glycerol stocks onto ½ YTSS agar plates containing antibiotics when required and incubated at 30 °C for 48 hours. A single colony was then used to inoculate a 10 mL ASW medium for a pre-culture, which was grown at 30 °C with shaking at 130 rpm for 48 hours to reach stationary phase. Cells were then diluted to an OD_600_ of 0.01 in either 75% algal spent medium or fresh ASW and supplemented, when indicated, with various concentrations of H_2_O_2_ (Bio-Lab ltd) or sterile water as control. Diluted cultures were transferred into a 96-well microtiter plate (150 µl per well) and overlaid with 50 µl hexadecane (Thermo Fisher Scientific) to prevent evaporation. Bacterial growth was monitored at 30°C using an Infinite 200 Pro M Plex plate reader (Tecan Group Ltd., Männedorf, Switzerland) with alternating cycles of 5 sec shaking and 59:55 min incubation. Absorbance at 600 nm was measured after each shaking cycle and multiplied by a factor of 3.86 to reflect optical density measurements performed in 1 cm cuvettes. Background absorbance, which is the measured OD_600_ of the culture media without bacterial addition, was subtracted from all measurements.

### Bacterial and algal growth during co-cultivation

For co-culture experiments, 30mL of ASW medium were inoculated with 10^4^ *E. huxleyi* cells, from late exponential phase inoculum, and incubated without shaking under the same conditions described above for algal cultures. After four days of algal growth, cultures were inoculated with 20 ul of *P. inhibens*, which had been pre-cultured for 48h in ASW, diluted to an OD_600_ of 0.01, and further diluted to a final concentration of 10^4^ cells/mL.

Bacterial growth was monitored at the indicated timepoints by serially diluting co-culture samples and plating them on 1⁄2 YTSS plates, with antibiotics when necessary. After two days of incubation at 30°C, colony-forming units (CFUs) were counted to calculate the total bacterial abundance.

Algal growth in cultures was monitored using a CellStream CS-100496 flow cytometer (Merck, Darmstadt, Germany), with excitation at 561nm and emission at 702 nm. For each sample, 30,000 events were recorded, and algal cells were gated based on event size and fluorescence intensity.

### Extracelullar H_2_O_2_ measurements

Extracellular H_2_O_2_ was quantified using the fluorescent probe Amplex red (Sigma-Aldrich), following the manufacturers instructions. Briefly, 100 μl of culture sample was centrifuged for 30 sec at 4000 rpm. Then, 10 μl of the supernatant was mixed with 40 μl of 1X Amplex reaction buffer in a μClear 96-well plate (Greiner Bio-One, Germany). Following that, 50 μl of 100 μM Amplex reagent was added to each well. The reaction was incubated in the dark for 30 min. Fluorescence was measured using an Infinite 200 Pro M Plex plate reader (Tecan Group Ltd., Männedorf, Switzerland) with excitation at 590nm and 530 nm emission. Background fluorescence was determined using control wells containing culture media, either minimal media or 75% spent media, without bacterial addition, and was subtracted from all individual measurements.

### H_2_O_2_ depletion by bacteria

Bacterial pre-cultures (prepared as described in *Bacterial growth curves* section) were diluted to an OD_600_ of 0.01 in 30 mL of fresh ASW medium. Cultures were supplemented with either 100 or 250 μM H_2_O_2_, or sterile water as control. Bacterial cultures were incubated at 30 °C with shaking at 130 rpm. Samples were collected at 0, 3, 6 and 24 h. H_2_O_2_ concentrations were measured as described in *Extracelullar H_2_O_2_ measurements* section. Bacterial growth (OD_600_) was monitored using an Ultrospec 2100 pro spectrophotometer (Biochrom, Cambridge, UK).

### Co-culture RNA-sequencing

The RNA-sequencing dataset used in this study was previously published by our group^23^. Briefly, co-cultures and pure cultures containing either *E. huxleyi* or *P. inhibens* were cultivated as described in *Bacterial and algal growth during co-cultivation* section. Cells were harvested by centrifugation at 4 °C, and total RNA was extracted using the ISOLATE II RNA Mini Kit (Meridian Bioscience, OH, USA). Ribosomal rRNA depletion was performed using Pan-Bacteria and custom probes designed for *E. huxleyi*. RNA libraries were deep sequenced on a NovaSeq 6000 system using a 100 cycles S2 Kit (Illumina, San Diego, CA, USA) in paired-end mode. Quality-filtered and trimmed reads were mapped to the *E. huxleyi* synthetic genome assembly (sGenome)^80^ or the *P. inhibens* DSM 17395 genome (accession: GCF_000154765.2). Raw sequencing data are available under BioProject accession PRJNA976961. Differential expression of algal genes in co-culture vs pure cultures was assessed using DESeq2. To compare absolute transcript abundances, feature counts were converted to transcripts per kilobase million (TPM), accounting for gene length and sequencing depth. Differential expression of bacterial genes was further analyzed using the limma-voom pipeline in R (v4.1.1) with the Bioconductor package limma^81^. Only genes with log₂ fold changes (logFC) and adjusted *p*-values <0.05 (Benjamini–Hochberg correction) were considered significant.

### P. inhibens pxn, katG, and sod null mutants

The primers and plasmids used for the KO constructs are described in table S5. DNA manipulation and cloning PCR was performed using Phusion High Fidelity DNA polymerase (Thermo Fisher Scientific), according with manufacturer recommended PCR conditions. PCR-amplified DNA was cleaned with NucleoSpin Gel and PCR Clean-up kit (MACHEREY-NAGEL, Düren, Germany).

For creation of *katg* null mutant (*Δkatg*) cells (ES329), ∼1000 bp regions upstream and downstream of the *katG* gene (accession: PGA1_RS19020) were amplified by PCR, using the primers 1-4. The gentamycin resistance marker of pBBR1MCS5^82^ was amplified using primers 13 and 14. The PCR-amplified fragments (upstream region + gentamycin resistance + downstream region) were assembled and cloned into the pCR™8/GW/TOPO® vector (Invitrogen, Thermo Fisher Scientific, Waltham, MA, USA) using restriction-free cloning^83^, generating the plasmid pDN22.

For creation of *pxn* null mutant (*Δpxn*) cells (ES151), ∼1000 bp regions upstream and downstream of the *pxn* gene (accession: PGA1_RS07285) were amplified by PCR, using the primers 7-10. The gentamycin resistance marker of pBBR1MCS5^82^ was amplified using primers 13 and 14. The PCR-amplified fragments (upstream region + gentamycin resistance + downstream region) were assembled and cloned pCR™8/GW/TOPO® vector (Invitrogen, Thermo Fisher Scientific, Waltham, MA, USA) using restriction-free cloning^83^, generating the plasmid pLY1.

For creation of *sod* null mutant (*Δsod*) cells (ES330), ∼1000 bp regions upstream and downstream of the *sod* gene (accession: PGA1_RS09485) were amplified by PCR, using the primers 17-20. The kanamycin resistance marker of pYDR1^19^ was amplified using primers 23 and 24. The PCR-amplified fragments (upstream region + kanamycin resistance + downstream region) were assembled and cloned into the pCR™8/GW/TOPO® vector (Invitrogen, Thermo Fisher Scientific, Waltham, MA, USA) using restriction-free cloning^83^, generating the plasmid pDN23.

*P. inhibens* electrocompetent cells (300 μl) were transformed with 10 μg of the constructed plasmids by a pulse of 2.5 kV (Bio Rad), and cells were selected on ½ YTSS plates containing 30 μg/ml Gentamycin or 150 μg/ml Kanamycin. Successful null mutants were verified in single cell clones by PCR (5-6 and 15-16 for *Δkatg*, 11-12 and 15-16 for *Δpxn,* and 21-22 and 25-26 for *Δsod*) and sanger sequencing.

### P. inhibens gene expression

Bacterial pre-cultures (prepared as described in *Bacterial growth curves* section) of *P. inhibens* were diluted to an OD_600_ of 0.01 in 30 mL of fresh ASW medium containing either 5.5 mM glucose or 6.6 mM betaine as the sole carbon source. Cultures were incubated at 30 °C with shaking at 130 rpm. After 22h cells were harvested by centrifugation at 4000 rpm for 10 min at 4°C. Cell pellets were resuspended in RNA lysis buffer (RLT, QIAGEN) containing 1% β-mercaptoethanol, transferred to tubes containing 300 mg of 100 µm acid-washed Low Binding Silica Beads (SPEX SamplePrep), and disrupted via bead beating at 30 mHz for 3 min. Total RNA was extracted using the Isolate II RNA Mini Kit (Meridian Bioscience, London, UK), followed by a second DNAse treatment using the TURBO DNase kit (Thermo Fisher Scientific) and further purified with RNAClean XP magnetic beads (Beckman Coulter). Total RNA was measured with the Qubit RNA HS Assay (Invitrogen, Thermo Fisher Scientific). RNA concentrations were quantified using the Quibit RNA HS Assay Kit (Invitrogen, Thermo Fisher Scientific). Equal amounts of RNA from each sample were used for cDNA synthesis with Superscript IV (Thermo Fisher Scientific). Quantitative PCR (qPCR) was performed in 384-well plates using the SensiFAST SYBR Lo-ROX Kit (Meridian Bioscience) on a QuantStudio 5 qPCR cycler (Applied Biosystems, Foster City, CA, USA). The program consisted of 40 amplification cycles following the enzyme requirements. Primers are listed in table S5. Efficiencies were determined using standard curves generated from serial dilutions of pooled cDNA. Only primer pairs with ≥85% efficiency were considered. Gene expression was calculated using the Comparative CT (ΔΔCT) method and normalized to the housekeeping genes *gyrA* and *recA*^22^.

### Betaine quantification from algal cultures

*E. huxleyi* cultures were grown as described in the *Strains and growth conditions* section. At defined timepoints (0, 4, 7, 10, 12 and 14 days), algal cell concentrations were quantified using a CellStream flow cytometer, as decribed in the *Bacterial and algal growth during co-cultivation* section. For each time point a full culture flask was used. For extracellular betaine measurements, 400 µl of culture were transferred to an eppendorf tube and centrifuged at 4000rpm for 5 min at 4°C. After centrifugation, 200 µl of the uppermost supernatant was transferred to a new tube and snap-frozen in liquid nitrogen. For intracellular betaine measurements, the remaining algal culture (∼29.450 mL) was centrifuged at 4000rpm for 5 min at 4°C. The supernatant was discarded, and the cell pellet was resuspended in 1mL of ASW, transferred to an eppendorf tube, and centrifuged again as described above. The supernatant was discarded using a pipette tip, and the pellet was snap-frozen in liquid nitrogen. All samples were stored at −80°C until for biological replicates were collected and processed together.

For metabolomics analysis, 750 µL of MeOH: DDW 7:3 containing 1.333 ug/ml ^13^C_5_^15^N-Betaine (Sigma-Aldrich) as internal standard, was added to the samples. After vortexing and three 10-minute sonication cycles, samples were centrifuged for 10 minutes at 14,000 rpm and 4°C, and the supernatant was collected. The pellets were re-extracted with 250 µL MeOH:DDW (7:3), followed by three 10-minute sonication cycles and centrifugation. Then the supernatants were combined with those from the first extraction, centrifuged again and transferred to vials. Samples with high concentration of betaine were diluted 20 times.

The measurement of betaine was performed as described at Zheng *et al*.^84^ with minor modifications described below. Briefly, analysis was performed using UPLC-ESI-MS/MS equipped with Acquity UPLC I class system (Waters). The LC separation was done using the Atlantis Premier BEH Z-HILIC (100 mm × 2.1 mm) (Waters). The Mobile phase B: acetonitrile and Mobile phase A: 20 mM ammonium carbonate with 0.1% ammonia hydroxide in water:acetonitrile (80:20, v/v). The flow rate was kept at 200 μl min−1 and gradient as follow: 0-2min 75% of B, 14 min 25% of B, 18 min 25% of B, 19 min 75% of B, for 4 min. Column temperature was set to 45°C and injection volume was 0.5 µl. MS detector (Waters TQ-XS) was equipped with ESI source. The source and de-solvation temperatures were maintained at 150°C and 600°C, respectively. The capillary voltage was set to 1.0 kV. Nitrogen was used as de-solvation gas and cone gas at the flow rate of 1000 L*h-1 and 150 L*h-1, respectively. Additional MS parameters are summarized in table S6. Data were processed with the MassLynx software with Targetlynx (Waters). Quantification of betaine was performed against an external calibration curve of Betaine (Sigma-Aldrich).

### Transcript abundance of H_2_O_2_ and betaine related genes in algal-rich ocean layers

Transcript abundances for orthologous groups (OG) of genes related to H_2_O_2_ and betaine metabolism were obtained from a previously published dataset from the TARA Oceans expedition^62^ and functionally annotated in the current study. Briefly, samples were collected from 126 globally distributed oceanic stations across different dephts. RNA was sequenced using an Illumina HiSeq2000 system in a paired-end mode. Sequencing reads were pre-procesed to generate High-quality (HQ) reads, which were then assembled into gene-encoding sequences to create the Ocean Microbial Reference Gene Catalog version 2 (OM-RGC.v2). Read counts were normalized by gene length and summarized based on EggNOG gene families (aligned to EggNOG version 3 database), and then divided by the transcript abundance of a constitutively expressed marker gene. DESeq2 was used for variance stabilization. The resulting transcript abundances are represented as log2-transformed values, indicating relative transcript numbers per cell.

We focused on the samples from light-penetrated, epipelagic zones, specifically surface waters (SRF), the deep chlorophyll maximum (DCM), and the mixed layer. Clorophyll a concentrations, used as a proxy for algal density, were obtained for each sample from the corresponding dataset (https://doi.org/10. 5281/zenodo.3473199) and included included in our analysis. To examine the relationship between gene expression and algal abundance, we used Pearson correlation analyses with the cor.test() function from the stats package in R. TPM-normalized gene expression values were correlated with chlorophyll *a* concentrations across samples. For each gene, we calculated a correlation coefficient (*r*) and corresponding *p*-value, which was adjusted for multiple testing using the Bonferroni method. Genes with adjusted *p*-values <0.05 were considered significantly correlated with chlorophyll *a*.

### H_2_O_2_-related genes co-expression analysis during a natural bloom

For the bloom expression analysis, we used a previously published metatranscriptomic dataset^64^, where transcriptomics reads were mapped to metagenome-assembled genomes (MAGs), and gene expression was normalized as transcripts per million (TPM). Relative expression of bacterial genes was assessed by assigning TPM values to specific MAGs and normalizing by genome size. The 46 most highly expressed MAGs were selected for further analysis, representing the following taxonomic groups: *Gammaproteobacteria* (36.95%)*, Bacteroidia* (32.6%)*, Alphaproteobacteria* (21.73%), *Acidimicrobiia* (4.34%), *Actinomycetia* (2.17%) and *Verrucomicrobiae* (2.17%).

We focused on the expression profiles of 21 genes related to H_2_O_2_ degradation and production, as well as betaine metabolism. Gene-level expression data were aggregated and summarized across 10 time points during the bloom, where chlorophyll *a*, representing algal abundance, was also measured.

To visualize temporal gene expression patterns, we generated a heatmap using Z-score normalized values. Z-scores were calculated by subtracting the mean and dividing by the standard deviation for each gene, enabling comparison of relative expression levels across time points.

## ACKNOWLEDGEMENTS

D.A.N.-B. received the Armando and Maria Jinich Fellowship. The study was funded by the Israel Science Foundation (ISF 692/24), the European Research Council (ERC StG 101075514) and the de Botton Center for Marine Science, granted to E.S.

## AUTHOR CONTRIBUTIONS

D.A.N.-B. and E.S. designed the study and wrote the manuscript. D.A.N.-B. and L.Y. connducted and analyzed the experimental work. D. Y. and V. L. analyzed the environmental data. S. M. performed the metabolomics analysis.

## COMPETING INTERESTS

The authors declare no competing interests.

## Notes

### Competing Interest Statement

The authors have declared no competing interest.

## REFERENCES

1. Morris, J.J., Rose, A.L., and Lu, Z. (2022). Reactive oxygen species in the world ocean and their impacts on marine ecosystems. Redox Biology 52, 102285. 10.1016/j.redox.2022.102285.

2. Lesser, M.P. (2006). OXIDATIVE STRESS IN MARINE ENVIRONMENTS: Biochemistry and Physiological Ecology. Annual Review of Physiology 68, 253–278. 10.1146/annurev.physiol.68.040104.110001.

3. Hopwood, M.J., Rapp, I., Schlosser, C., and Achterberg, E.P. (2017). Hydrogen peroxide in deep waters from the Mediterranean Sea, South Atlantic and South Pacific Oceans. Sci Rep 7, 43436. 10.1038/srep43436.

4. Obernosterer, I., Ruardij, P., and Herndl, G.J. (2001). Spatial and diurnal dynamics of dissolved organic matter (DOM) fluorescence and H2O2 and the photochemical oxygen demand of surface water DOM across the subtropical Atlantic Ocean. Limnology and Oceanography 46, 632–643. 10.4319/lo.2001.46.3.0632.

5. Cooper, W.J., Saltzman, E.S., and Zika, R.G. (1987). The contribution of rainwater to variability in surface ocean hydrogen peroxide. Journal of Geophysical Research: Oceans 92, 2970–2980. 10.1029/JC092iC03p02970.

6. Zinser, E.R. (2018). The microbial contribution to reactive oxygen species dynamics in marine ecosystems. Environmental Microbiology Reports 10, 412–427. 10.1111/1758-2229.12626.

7. Diaz, J.M., and Plummer, S. (2018). Production of extracellular reactive oxygen species by phytoplankton: past and future directions. J Plankton Res 40, 655–666. 10.1093/plankt/fby039.

8. Hansel, C.M., and Diaz, J.M. (2021). Production of Extracellular Reactive Oxygen Species by Marine Biota. Annual Review of Marine Science 13, 177–200. 10.1146/annurev-marine-041320-102550.

9. Bond, R.J., Hansel, C.M., and Voelker, B.M. (2020). Heterotrophic Bacteria Exhibit a Wide Range of Rates of Extracellular Production and Decay of Hydrogen Peroxide. Frontiers in Marine Science 7.

10. Mishra, S., and Imlay, J. (2012). Why do bacteria use so many enzymes to scavenge hydrogen peroxide? Arch Biochem Biophys 525, 145–160. 10.1016/j.abb.2012.04.014.

11. Zinser, E.R. (2018). Cross-protection from hydrogen peroxide by helper microbes: the impacts on the cyanobacterium Prochlorococcus and other beneficiaries in marine communities. Environmental Microbiology Reports 10, 399–411. 10.1111/1758-2229.12625.

12. Omar, N.M., Prášil, O., McCain, J.S.P., and Campbell, D.A. (2022). Diffusional Interactions among Marine Phytoplankton and Bacterioplankton: Modelling H2O2 as a Case Study. Microorganisms 10, 821. 10.3390/microorganisms10040821.

13. Varaljay, V.A., Robidart, J., Preston, C.M., Gifford, S.M., Durham, B.P., Burns, A.S., Ryan, J.P., Marin III, R., Kiene, R.P., Zehr, J.P., et al. (2015). Single-taxon field measurements of bacterial gene regulation controlling DMSP fate. ISME J 9, 1677– 1686. 10.1038/ismej.2015.23.

14. Bendif, E.M., Nevado, B., Wong, E.L.Y., Hagino, K., Probert, I., Young, J.R., Rickaby, R.E.M., and Filatov, D.A. (2019). Repeated species radiations in the recent evolution of the key marine phytoplankton lineage Gephyrocapsa. Nature Communications 10, 4234. 10.1038/s41467-019-12169-7.

15. Deng, Y., Yu, R., Grabe, V., Sommermann, T., Werner, M., Vallet, M., Zerfaß, C., Werz, O., and Pohnert, G. (2024). Bacteria modulate microalgal aging physiology through the induction of extracellular vesicle production to remove harmful metabolites. Nat Microbiol 9, 2356–2368. 10.1038/s41564-024-01746-2.

16. Sheyn, U., Rosenwasser, S., Ben-Dor, S., Porat, Z., and Vardi, A. (2016). Modulation of host ROS metabolism is essential for viral infection of a bloom-forming coccolithophore in the ocean. ISME J 10, 1742–1754. 10.1038/ismej.2015.228.

17. Segev, E., Wyche, T.P., Kim, K.H., Petersen, J., Ellebrandt, C., Vlamakis, H., Barteneva, N., Paulson, J.N., Chai, L., Clardy, J., et al. (2016). Dynamic metabolic exchange governs a marine algal-bacterial interaction. eLife 5, e17473. 10.7554/eLife.17473.

18. Seyedsayamdost, M.R., Case, R.J., Kolter, R., and Clardy, J. (2011). The Jekyll-and-Hyde chemistry of *Phaeobacter gallaeciensis*. Nature Chem 3, 331–335. 10.1038/nchem.1002.

19. Abada, A., Beiralas, R., Narvaez, D., Sperfeld, M., Duchin-Rapp, Y., Lipsman, V., Yuda, L., Cohen, B., Carmieli, R., Ben-Dor, S., et al. (2023). Aerobic bacteria produce nitric oxide via denitrification and promote algal population collapse. ISME J, 1–17. 10.1038/s41396-023-01427-8.

20. Beiralas, R., Ozer, N., and Segev, E. (2023). Abundant Sulfitobacter marine bacteria protect Emiliania huxleyi algae from pathogenic bacteria. ISME COMMUN. 3, 1–10. 10.1038/s43705-023-00311-y.

21. Majzoub, M.E., Beyersmann, P.G., Simon, M., Thomas, T., Brinkhoff, T., and Egan, S. (2019). Phaeobacter inhibens controls bacterial community assembly on a marine diatom. FEMS Microbiology Ecology 95, fiz060. 10.1093/femsec/fiz060.

22. Lipsman, V., Shlakhter, O., Rocha, J., and Segev, E. (2024). Bacteria contribute exopolysaccharides to an algal-bacterial joint extracellular matrix. npj Biofilms Microbiomes 10, 1–16. 10.1038/s41522-024-00510-y.

23. Sperfeld, M., Narváez-Barragán, D.A., Malitsky, S., Frydman, V., Yuda, L., Rocha, J., and Segev, E. (2024). Algal methylated compounds shorten the lag phase of Phaeobacter inhibens bacteria. Nat Microbiol, 1–16. 10.1038/s41564-024-01742-6.

24. Bramucci, A.R., and Case, R.J. (2019). Phaeobacter inhibens induces apoptosis-like programmed cell death in calcifying Emiliania huxleyi. Sci Rep 9, 5215. 10.1038/s41598-018-36847-6.

25. Wang, R., Gallant, É., Wilson, M.Z., Wu, Y., Li, A., Gitai, Z., and Seyedsayamdost, M.R. (2022). Algal p-coumaric acid induces oxidative stress and siderophore biosynthesis in the bacterial symbiont Phaeobacter inhibens. Cell Chemical Biology 29, 670–679.e5. 10.1016/j.chembiol.2021.08.002.

26. Dunand, C., Crèvecoeur, M., and Penel, C. (2007). Distribution of superoxide and hydrogen peroxide in Arabidopsis root and their influence on root development: possible interaction with peroxidases. New Phytologist 174, 332–341. 10.1111/j.1469-8137.2007.01995.x.

27. Strom, S.L., Bright, K.J., Fredrickson, K.A., and Cooney, E.C. (2018). Phytoplankton defenses: Do Emiliania huxleyi coccoliths protect against microzooplankton predators? Limnology and Oceanography 63, 617–627. 10.1002/lno.10655.

28. Plummer, S., Taylor, A.E., Harvey, E.L., Hansel, C.M., and Diaz, J.M. (2019). Dynamic Regulation of Extracellular Superoxide Production by the Coccolithophore Emiliania huxleyi (CCMP 374). Front. Microbiol. 10. 10.3389/fmicb.2019.01546.

29. Hancock, T.L., Dahedl, E.K., Kratz, M.A., and Urakawa, H. (2024). The synchronicity of bloom-forming cyanobacteria transcription patterns and hydrogen peroxide dynamics. Environmental Pollution 348, 123812. 10.1016/j.envpol.2024.123812.

30. Pospíšil, P., Kumar, A., and Prasad, A. (2022). Reactive oxygen species in photosystem II: relevance for oxidative signaling. Photosynth Res 152, 245–260. 10.1007/s11120-022-00922-x.

31. Kim, K., and Portis, A.R. (2004). Oxygen-dependent H2O2 production by Rubisco. FEBS Letters 571, 124–128. 10.1016/j.febslet.2004.06.064.

32. Nikkanen, L., Solymosi, D., Jokel, M., and Allahverdiyeva, Y. (2021). Regulatory electron transport pathways of photosynthesis in cyanobacteria and microalgae: Recent advances and biotechnological prospects. Physiologia Plantarum 173, 514–525. 10.1111/ppl.13404.

33. Caspi, R., Billington, R., Fulcher, C.A., Keseler, I.M., Kothari, A., Krummenacker, M., Latendresse, M., Midford, P.E., Ong, Q., Ong, W.K., et al. (2018). The MetaCyc database of metabolic pathways and enzymes. Nucleic Acids Research 46, D633– D639. 10.1093/nar/gkx935.

34. Sandamalika, W.M.G., Udayantha, H.M.V., Liyanage, D.S., Lim, C., Kim, G., Kwon, H., and Lee, J. (2022). Identification of reactive oxygen species modulator 1 (Romo 1) from black rockfish (*Sebastes schlegelii*) and deciphering its molecular characteristics, immune responses, oxidative stress modulation, and wound healing properties. Fish & Shellfish Immunology 125, 266–275. 10.1016/j.fsi.2022.05.026.

35. Shin, J.A., Chung, J.S., Cho, S.-H., Kim, H.J., and Yoo, Y.D. (2013). Romo1 expression contributes to oxidative stress-induced death of lung epithelial cells. Biochemical and Biophysical Research Communications 439, 315–320. 10.1016/j.bbrc.2013.07.012.

36. Chaturvedi, R., Cheng, Y., Asim, M., Bussière, F.I., Xu, H., Gobert, A.P., Hacker, A., Casero, R.A., and Wilson, K.T. (2004). Induction of Polyamine Oxidase 1 by *Helicobacter pylori* Causes Macrophage Apoptosis by Hydrogen Peroxide Release and Mitochondrial Membrane Depolarization*. Journal of Biological Chemistry 279, 40161–40173. 10.1074/jbc.M401370200.

37. Wittke, I., Wiedemeyer, R., Pillmann, A., Savelyeva, L., Westermann, F., and Schwab, M. (2003). Neuroblastoma-Derived Sulfhydryl Oxidase, a New Member of the Sulfhydryl Oxidase/Quiescin6 Family, Regulates Sensitization to Interferon γ-Induced Cell Death in Human Neuroblastoma Cells. Cancer Research 63, 7742–7752.

38. Zhang, H., Wang, H., Zheng, W., Yao, Z., Peng, Y., Zhang, S., Hu, Z., Tao, Z., and Zheng, T. (2017). Toxic Effects of Prodigiosin Secreted by Hahella sp. KA22 on Harmful Alga Phaeocystis globosa. Front. Microbiol. 8. 10.3389/fmicb.2017.00999.

39. Rojas, C.M., Senthil-Kumar, M., Wang, K., Ryu, C.-M., Kaundal, A., and Mysore, K.S. (2012). Glycolate Oxidase Modulates Reactive Oxygen Species–Mediated Signal Transduction during Nonhost Resistance in *Nicotiana benthamiana* and *Arabidopsis*. The Plant Cell 24, 336–352. 10.1105/tpc.111.093245.

40. Ferrer, M.D., Tauler, P., Sureda, A., Palacín, C., Tur, J.A., and Pons, A. (2013). Antioxidants restore protoporphyrinogen oxidase in variegate porphyria patients. European Journal of Clinical Investigation 43, 668–678. 10.1111/eci.12091.

41. Sen, A., and Imlay, J.A. (2021). How Microbes Defend Themselves From Incoming Hydrogen Peroxide. Front. Immunol. 12. 10.3389/fimmu.2021.667343.

42. Martins, D., Kathiresan, M., and English, A.M. (2013). Cytochrome *c* peroxidase is a mitochondrial heme-based H2O2 sensor that modulates antioxidant defense. Free Radical Biology and Medicine 65, 541–551. 10.1016/j.freeradbiomed.2013.06.037.

43. Dubbs, J.M., and Mongkolsuk, S. (2007). Peroxiredoxins in Bacterial Antioxidant Defense. In Peroxiredoxin Systems: Structures and Functions, L. Flohé and J. R. Harris, eds. (Springer Netherlands), pp. 143–193. 10.1007/978-1-4020-6051-9_7.

44. Storz, G., and Tartaglia, L.A. (1992). OxyR: a regulator of antioxidant genes. J Nutr 122, 627–630. 10.1093/jn/122.suppl_3.627.

45. Seaver, L.C., and Imlay, J.A. (2001). Hydrogen Peroxide Fluxes and Compartmentalization inside Growing Escherichia coli. Journal of Bacteriology 183, 7182–7189. 10.1128/jb.183.24.7182-7189.2001.

46. Smith, D.J., Berry, M.A., Cory, R.M., Johengen, T.H., Kling, G.W., Davis, T.W., and Dick, G.J. (2022). Heterotrophic Bacteria Dominate Catalase Expression during Microcystis Blooms. Appl Environ Microbiol 88, e02544–21. 10.1128/aem.02544-21.

47. Hébrard, M., Viala, J.P.M., Méresse, S., Barras, F., and Aussel, L. (2009). Redundant Hydrogen Peroxide Scavengers Contribute to Salmonella Virulence and Oxidative Stress Resistance. Journal of Bacteriology 191, 4605–4614. 10.1128/jb.00144-09.

48. Wang, Y., Branicky, R., Noë, A., and Hekimi, S. (2018). Superoxide dismutases: Dual roles in controlling ROS damage and regulating ROS signaling. J Cell Biol 217, 1915– 1928. 10.1083/jcb.201708007.

49. Hansel, C.M., Diaz, J.M., and Plummer, S. (2019). Tight Regulation of Extracellular Superoxide Points to Its Vital Role in the Physiology of the Globally Relevant Roseobacter Clade. mBio 10, 10.1128/mbio.02668-18. 10.1128/mbio.02668-18.

50. Moran, M.A., Reisch, C.R., Kiene, R.P., and Whitman, W.B. (2012). Genomic insights into bacterial DMSP transformations. Annu. Rev. Mar. Sci. 4, 523–542. 10.1146/annurev-marine-120710-100827.

51. Lahham, M., Jha, S., Goj, D., Macheroux, P., and Wallner, S. (2021). The family of sarcosine oxidases: Same reaction, different products. Archives of Biochemistry and Biophysics 704, 108868. 10.1016/j.abb.2021.108868.

52. Karp, P.D., Billington, R., Caspi, R., Fulcher, C.A., Latendresse, M., Kothari, A., Keseler, I.M., Krummenacker, M., Midford, P.E., Ong, Q., et al. (2019). The BioCyc collection of microbial genomes and metabolic pathways. Briefings in Bioinformatics 20, 1085–1093. 10.1093/bib/bbx085.

53. Mausz, M.A., Airs, R.L., Dixon, J.L., Widdicombe, C.E., Tarran, G.A., Polimene, L., Dashfield, S., Beale, R., Scanlan, D.J., and Chen, Y. (2022). Microbial uptake dynamics of choline and glycine betaine in coastal seawater. Limnology and Oceanography 67, 1052–1064. 10.1002/lno.12056.

54. Thole, S., Kalhoefer, D., Voget, S., Berger, M., Engelhardt, T., Liesegang, H., Wollherr, A., Kjelleberg, S., Daniel, R., Simon, M., et al. (2012). Phaeobacter gallaeciensis genomes from globally opposite locations reveal high similarity of adaptation to surface life. ISME J 6, 2229–2244. 10.1038/ismej.2012.62.

55. Narváez-Barragán, D.A., Sperfeld, M., and Segev, E. DmdA-independent lag phase shortening in Phaeobacter inhibens bacteria under stress conditions. The FEBS Journal n/a. 10.1111/febs.70128.

56. Wu, H., Wang, Y., Ying, M., Jin, C., Li, J., and Hu, X. (2021). Lactate dehydrogenases amplify reactive oxygen species in cancer cells in response to oxidative stimuli. Signal Transduction and Targeted Therapy 6, 242. 10.1038/s41392-021-00595-3.

57. di Salvo, M.L., Safo, M.K., Musayev, F.N., Bossa, F., and Schirch, V. (2003). Structure and mechanism of *Escherichia coli* pyridoxine 5′-phosphate oxidase. Biochimica et Biophysica Acta (BBA) - Proteins and Proteomics 1647, 76–82. 10.1016/S1570-9639(03)00060-8.

58. Skotnicová, P., Sobotka, R., Shepherd, M., Hájek, J., Hrouzek, P., and Tichý, M. (2018). The cyanobacterial protoporphyrinogen oxidase HemJ is a new b-type heme protein functionally coupled with coproporphyrinogen III oxidase. Journal of Biological Chemistry 293, 12394–12404. 10.1074/jbc.RA118.003441.

59. Zeng, J., Sun, Q., Su, J., Han, J., Zhang, Q., and Jin, Y. (2015). Protoporphyrin IX catalyzed hydrogen peroxide to generate singlet oxygen. International Journal of Clinical and Experimental Medicine 8, 6829.

60. Zecher, K., Hayes, K.R., and Philipp, B. (2020). Evidence of Interdomain Ammonium Cross-Feeding From Methylamine- and Glycine Betaine-Degrading Rhodobacteraceae to Diatoms as a Widespread Interaction in the Marine Phycosphere. Front. Microbiol. 11. 10.3389/fmicb.2020.533894.

61. Boysen, A.K., Durham, B.P., Kumler, W., Key, R.S., Heal, K.R., Carlson, L.T., Groussman, R.D., Armbrust, E.V., and Ingalls, A.E. (2022). Glycine betaine uptake and metabolism in marine microbial communities. Environmental Microbiology 24, 2380–2403. 10.1111/1462-2920.16020.

62. Salazar, G., Paoli, L., Alberti, A., Huerta-Cepas, J., Ruscheweyh, H.-J., Cuenca, M., Field, C.M., Coelho, L.P., Cruaud, C., Engelen, S., et al. (2019). Gene Expression Changes and Community Turnover Differentially Shape the Global Ocean Metatranscriptome. Cell 179, 1068–1083.e21. 10.1016/j.cell.2019.10.014.

63. Tyrrell, T., and Merico, A. (2004). Emiliania huxleyi: bloom observations and the conditions that induce them. In Coccolithophores: From Molecular Processes to Global Impact, H. R. Thierstein and J. R. Young, eds. (Springer), pp. 75–97. 10.1007/978-3-662-06278-4_4.

64. Sidhu, C., Kirstein, I.V., Meunier, C.L., Rick, J., Fofonova, V., Wiltshire, K.H., Steinke, N., Vidal-Melgosa, S., Hehemann, J.-H., Huettel, B., et al. (2023). Dissolved storage glycans shaped the community composition of abundant bacterioplankton clades during a North Sea spring phytoplankton bloom. Microbiome 11, 77. 10.1186/s40168-023-01517-x.

65. Ticak, T., Kountz, D.J., Girosky, K.E., Krzycki, J.A., and Ferguson, D.J. (2014). A nonpyrrolysine member of the widely distributed trimethylamine methyltransferase family is a glycine betaine methyltransferase. Proceedings of the National Academy of Sciences 111, E4668–E4676. 10.1073/pnas.1409642111.

66. Wargo, M.J., Szwergold, B.S., and Hogan, D.A. (2008). Identification of Two Gene Clusters and a Transcriptional Regulator Required for Pseudomonas aeruginosa Glycine Betaine Catabolism. Journal of Bacteriology 190, 2690–2699. 10.1128/jb.01393-07.

67. Seymour, J.R., Amin, S.A., Raina, J.-B., and Stocker, R. (2017). Zooming in on the phycosphere: the ecological interface for phytoplankton–bacteria relationships. Nat Microbiol 2, 1–12. 10.1038/nmicrobiol.2017.65.

68. Kim, S., Bang, Y.-J., Kim, D., Lim, J.G., Oh, M.H., and Choi, S.H. (2014). Distinct characteristics of OxyR2, a new OxyR-type regulator, ensuring expression of Peroxiredoxin 2 detoxifying low levels of hydrogen peroxide in ibrio vulnificus. Molecular Microbiology 93, 992–1009. 10.1111/mmi.12712.

69. Seaver, L.C., and Imlay, J.A. (2001). Alkyl Hydroperoxide Reductase Is the Primary Scavenger of Endogenous Hydrogen Peroxide in Escherichia coli. Journal of Bacteriology 183, 7173–7181. 10.1128/jb.183.24.7173-7181.2001.

70. Chen, C., Wang, Y., Chen, K., Shi, X., and Yang, G. (2021). Using hydrogen peroxide to control cyanobacterial blooms: A mesocosm study focused on the effects of algal density in Lake Chaohu, China. Environmental Pollution 272, 115923. 10.1016/j.envpol.2020.115923.

71. Machado, M.D., and Soares, E.V. (2022). Life and death of Pseudokirchneriella subcapitata: physiological changes during chronological aging. Appl Microbiol Biotechnol 106, 8245–8258. 10.1007/s00253-022-12267-5.

72. Carlioz, A., and Touati, D. (1986). Isolation of superoxide dismutase mutants in Escherichia coli: is superoxide dismutase necessary for aerobic life? The EMBO Journal 5, 623–630. 10.1002/j.1460-2075.1986.tb04256.x.

73. Storz, G., Christman, M.F., Sies, H., and Ames, B.N. (1987). Spontaneous mutagenesis and oxidative damage to DNA in Salmonella typhimurium. Proceedings of the National Academy of Sciences 84, 8917–8921. 10.1073/pnas.84.24.8917.

74. Vardi, A., Formiggini, F., Casotti, R., Martino, A.D., Ribalet, F., Miralto, A., and Bowler, C. (2006). A Stress Surveillance System Based on Calcium and Nitric Oxide in Marine Diatoms. PLOS Biology 4, e60. 10.1371/journal.pbio.0040060.

75. Niu, L., and Liao, W. (2016). Hydrogen Peroxide Signaling in Plant Development and Abiotic Responses: Crosstalk with Nitric Oxide and Calcium. Front. Plant Sci. 7. 10.3389/fpls.2016.00230.

76. Barak-Gavish, N., Dassa, B., Kuhlisch, C., Nussbaum, I., Brandis, A., Rosenberg, G., Avraham, R., and Vardi, A. (2023). Bacterial lifestyle switch in response to algal metabolites. eLife 12, e84400. 10.7554/eLife.84400.

77. Raina, J.-B., Lambert, B.S., Parks, D.H., Rinke, C., Siboni, N., Bramucci, A., Ostrowski, M., Signal, B., Lutz, A., Mendis, H., et al. (2022). Chemotaxis shapes the microscale organization of the ocean’s microbiome. Nature 605, 132–138. 10.1038/s41586-022-04614-3.

78. Keller, M.D., Kiene, R.P., Matrai, P.A., and Bellows, W.K. (1999). Production of glycine betaine and dimethylsulfoniopropionate in marine phytoplankton. I. Batch cultures. Marine Biology 135, 237–248. 10.1007/s002270050621.

79. Goyet, C., and Poisson, A. (1989). New determination of carbonic acid dissociation constants in seawater as a function of temperature and salinity. Deep Sea Research Part A. Oceanographic Research Papers 36, 1635–1654. 10.1016/0198-0149(89)90064-2.

80. Sperfeld Martin, Yahalomi Dayana, and Segev Einat (2022). Resolving the Microalgal Gene Landscape at the Strain Level: a Novel Hybrid Transcriptome of Emiliania huxleyi CCMP3266. Applied and Environmental Microbiology 88, e01418–21. 10.1128/AEM.01418-21.

81. Ritchie, M.E., Phipson, B., Wu, D., Hu, Y., Law, C.W., Shi, W., and Smyth, G.K. (2015). limma powers differential expression analyses for RNA-sequencing and microarray studies. Nucleic Acids Research 43, e47. 10.1093/nar/gkv007.

82. Kovach, M.E., Elzer, P.H., Steven Hill, D., Robertson, G.T., Farris, M.A., Roop, R.M., and Peterson, K.M. (1995). Four new derivatives of the broad-host-range cloning vector pBBR1MCS, carrying different antibiotic-resistance cassettes. Gene 166, 175–176. 10.1016/0378-1119(95)00584-1.

83. Peleg, Y., and Unger, T. (2014). Application of the Restriction-Free (RF) cloning for multicomponents assembly. In DNA Cloning and Assembly Methods Methods in Molecular Biology., S. Valla and R. Lale, eds. (Humana Press), pp. 73–87. 10.1007/978-1-62703-764-8_6.

84. Zheng, L., Cardaci, S., Jerby, L., MacKenzie, E.D., Sciacovelli, M., Johnson, T.I., Gaude, E., King, A., Leach, J.D.G., Edrada-Ebel, R., et al. (2015). Fumarate induces redox-dependent senescence by modifying glutathione metabolism. Nat Commun 6, 6001. 10.1038/ncomms7001.

