## Supplementary Information for "Algal Betaine Triggers Bacterial Hydrogen Peroxide (H_2_O_2_) Production that Promotes Algal Demise"

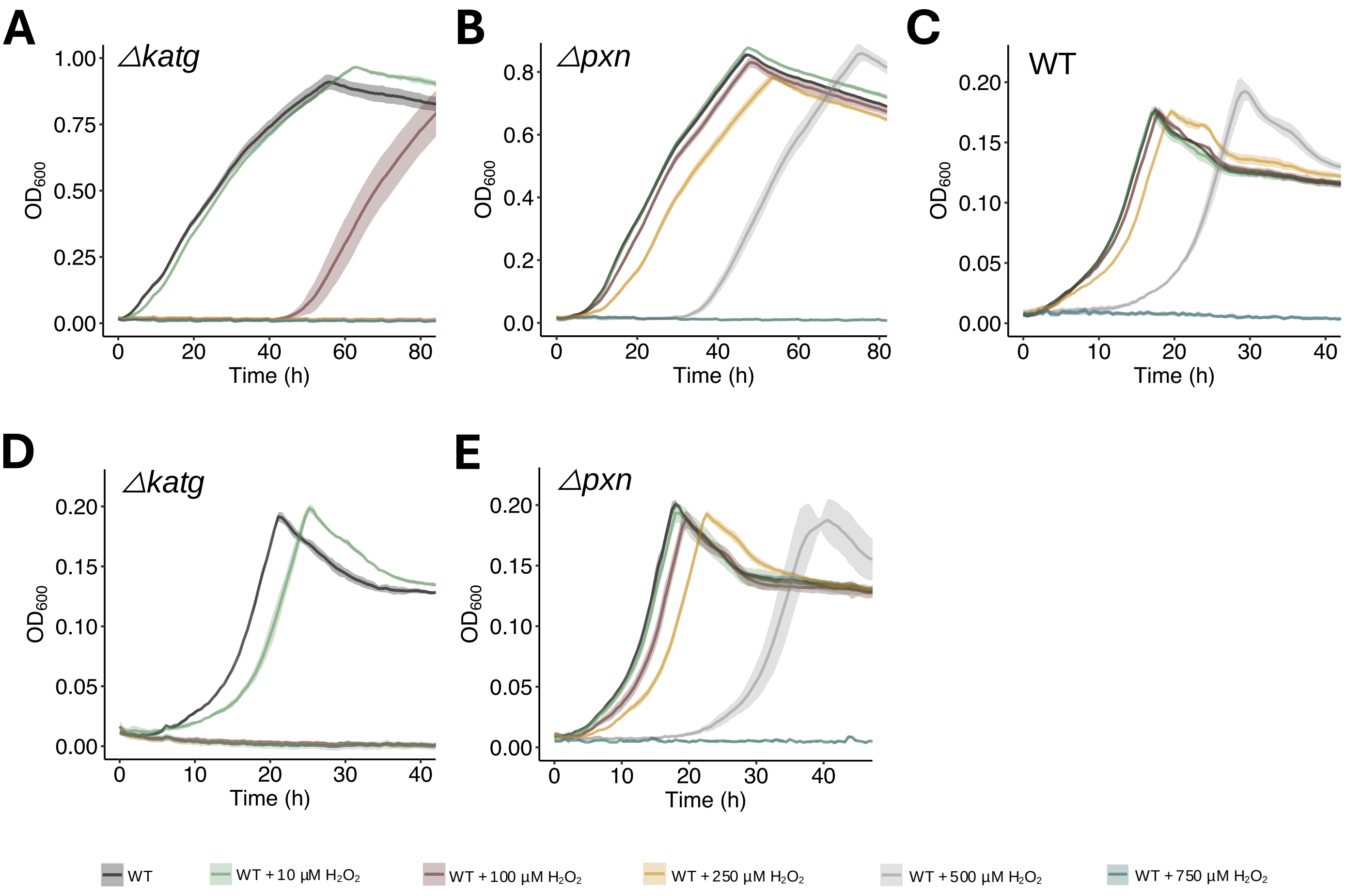


**S1. $H_2O_2$ susceptibility of *P. inhibens* mutants impaired in $H_2O_2$ degradation. A)** *Δkatg* and **B)** *Δpxn* bacteria grown in minimal medium with 5.5 mM glucose as a carbon source. **C)** WT, **D)** *Δkatg* and **E)** *Δpxn* bacteria grown in minimal medium with 1 mM glucose as a carbon source. All cultures were supplemented with 0 (black), 10 (green), 100 (brown), 250 (yellow), 500 (gray), and 750 (blue) µM $H_2O_2$. Lines represent the average growth curve based on three biological replicates and shaded areas indicate the standard deviation (SD)

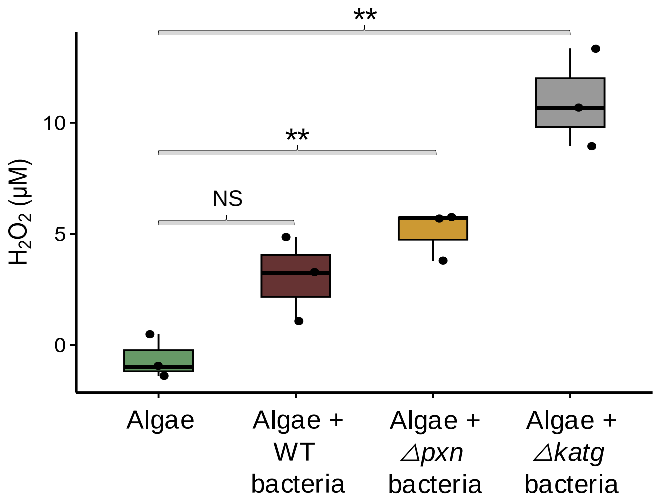


**S2. Accumulation of $H_2O_2$ during algal demise in co-cultures with *Δpxn* and *Δkatg* bacterial mutants.** Extracellular $H_2O_2$ concentration (µM) measured at day 12, in cultures of *E. huxleyi* grown alongside WT (brown), *Δpxn* (yellow), or *ΔkatG* (gray) bacteria. Box-plots show results from three independent experiments; black dots indicate individual measurements. Box-plot elements: center line - median; box limits -upper and lower quartiles; whiskers – min and max values. Normality was first assessed using a Shapiro–Wilk test, followed by an unpaired t-test to determine statistical differences between mono- and co-cultures at various time points. NS: Not significant,*P≤0.002.

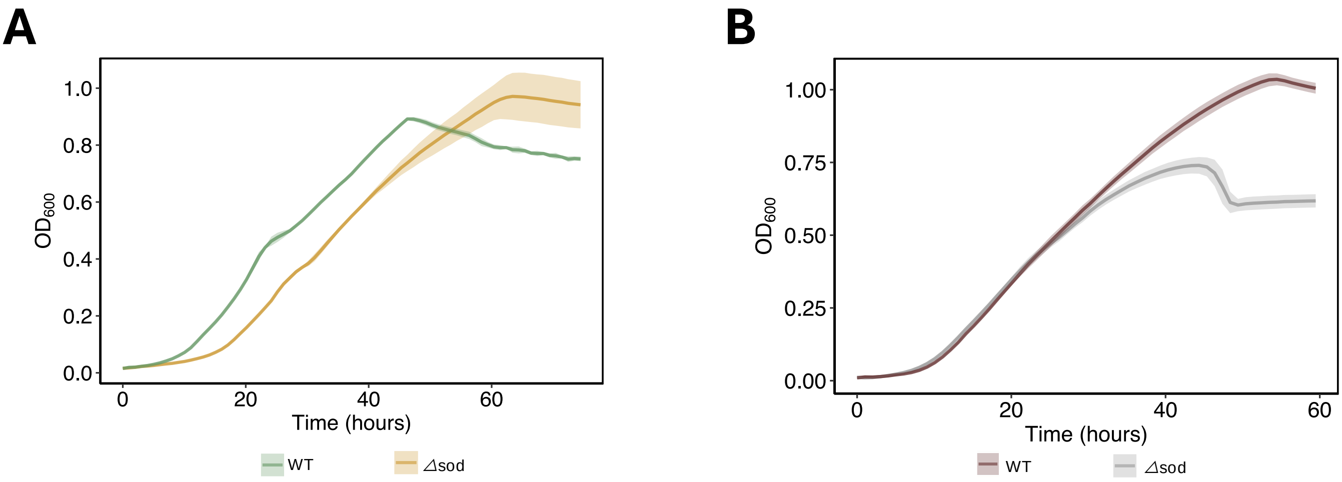


**S3. Growth of the mutant Δ*sod* compared with WT bacteria.** Bacteria were cultured in **A)** minimal medium with 5.5mM glucose as a carbon source (WT: green and *Δsod* mutant: yellow), or **B)** 75% spent media collected from algae at day 14 (WT: Brown and *Δsod:* gray). Lines represent the average growth curve based on three biological replicates and shaded areas indicate the standard deviation (SD).

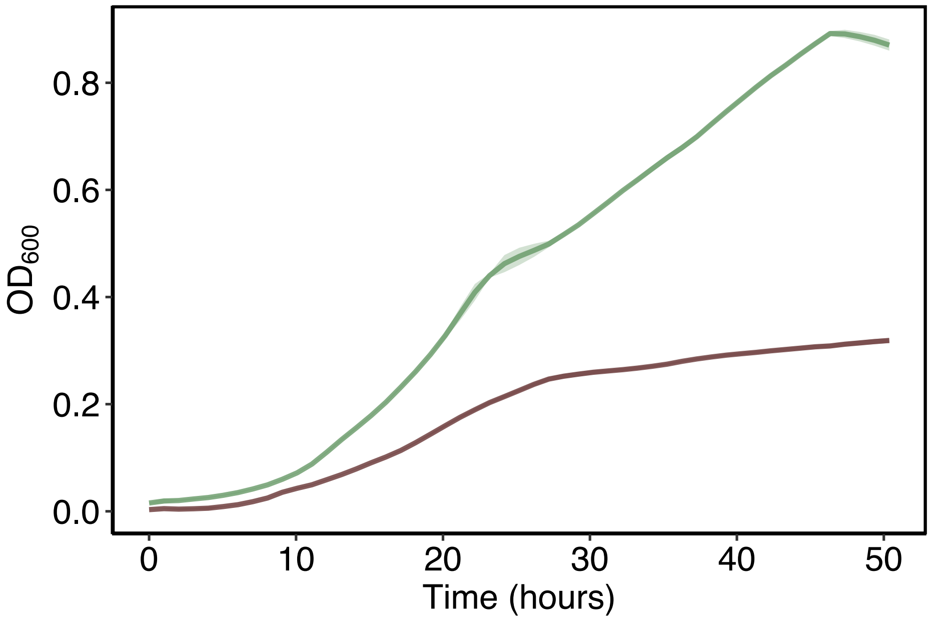


**S4. *P. inhibens* bacteria can utilize betaine as a sole carbon source.** Bacterial growth in ASW medium as described in the Material and methods section '*Media and strains'* utilizing 6.6mM betaine (brown) or 5.5mM glucose (green) as a sole carbon source. Lines represent the average growth curve based on three biological replicates and shaded areas indicate the standard deviation (SD).

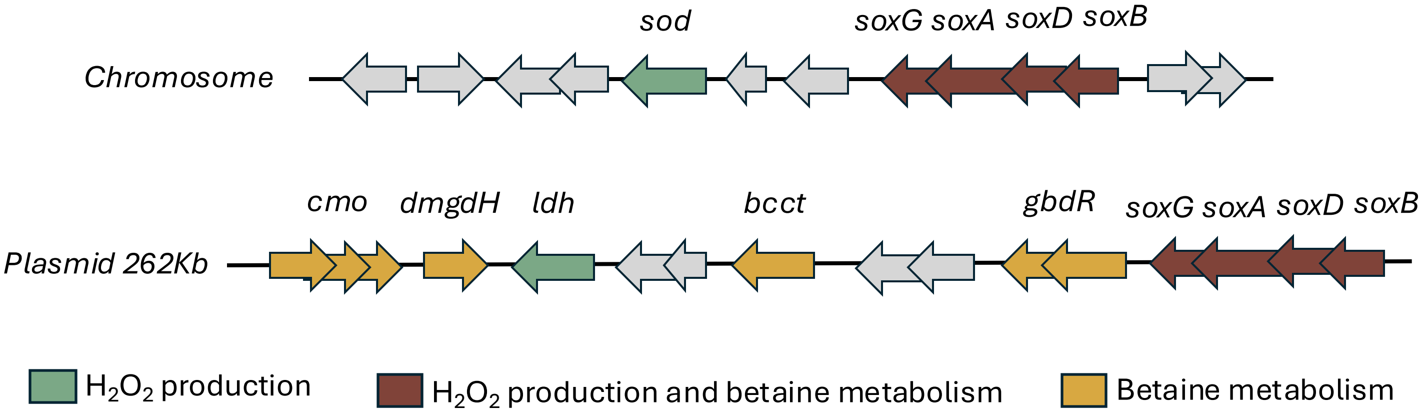


**S5. Bacterial genetic potential of betaine metabolism and $H_2O_2$ production.** Genomic context of bacterial genes related to betaine metabolism (yellow), $H_2O_2$ production (green), or both (Brown) on the *P. inhibens* chromosome and 262Kb plasmid.

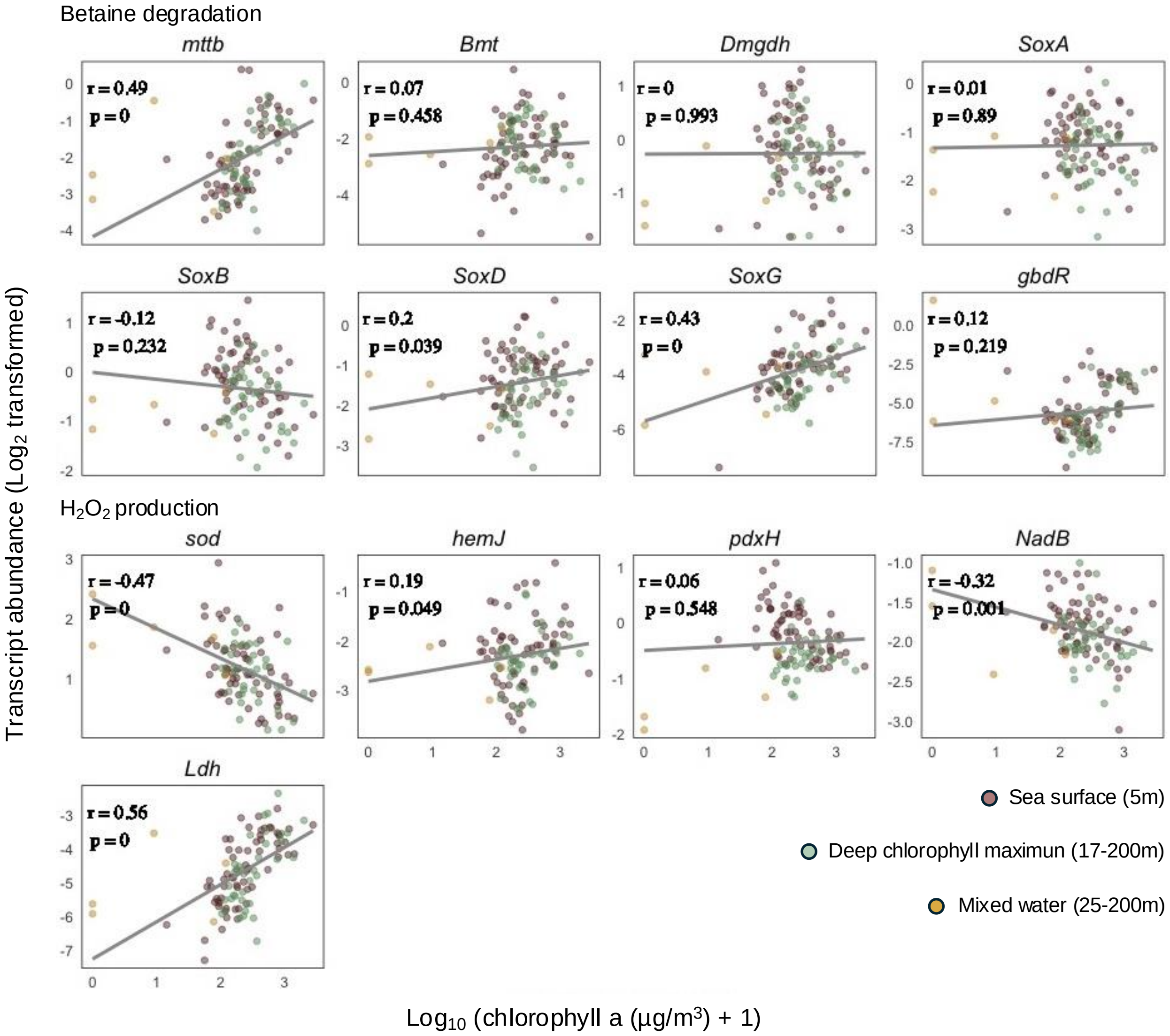


**S6. Transcript abundance related to betaine metabolism and $H_2O_2$-production in algal-rich layers of the marine environment.** Transcript abundances of orthologous groups (OG) of genes encoding proteins involved in betaine metabolism and $H_2O_2$ production in the epipelagic region, collected by the TARA expedition. The epipelagic region consists of the sea surface (brown), deep chlorophyll maximum (green) and mixed water layer samples (yellow). Transcript abundances (Log2 transformed) were correlated with chlorophyll a concentrations (Log10 (µg/m3 + 1) transformed) at the sampling points. r: Spearman's correlation. p: *p*-value adjusted using the Bonferroni correction. Data adapted from Salazar *et al.* [2].

**Table S1. Algal genes involved in $H_2O_2$ production and degradation.** Log2 fold changes (logFC) in *E. huxleyi* gene expression during co-cultivation with bacteria (A+B), compared to axenic cultures (A) across different growth phases. The demise phase in co-cultures corresponds to the late stationary phase in axenic algal cultures. padj refers to the adjusted *p*-value calculated using the Bonferroni correction. The underlying dataset was previously described in Sperfeld & Narváez-Barragán *et al*.[1]

| | | | | A+B vs A early exponential | | A+B vs A late exponential | | A+B vs A stationary | | A+B vs A demise | |
|---|---|---|---|---|---|---|---|---|---|---|---|
| **Function** | **Gene ID** | **Annotation** | **Short name** | **logFC** | **padj** | **logFC** | **padj** | **logFC** | **padj** | **logFC** | **padj** |
| $H_2O_2$ production | G20714 | reactive oxygen species modulator 1 | romo1 | 0.002 | 0.987 | -0.032 | 0.884 | -0.025 | 0.333 | 1.979 | 0.000 |
| | G21128 | Superoxide dismutase | sod1 | 0.014 | 0.934 | -0.676 | 0.024 | -0.005 | 0.963 | 3.584 | 0.000 |
| | G20784 | superoxide dismutase, Cu-Zn | sod2 | -0.004 | NA | -0.028 | 0.831 | -0.010 | 0.774 | 1.549 | 0.001 |
| | G3866 | Cu/Zn superoxide dismutase | sod3 | 0.007 | NA | -0.004 | 0.977 | -0.001 | 0.992 | -0.053 | 0.923 |
| | G16991 | Superoxide dismutase | sod4 | 0.021 | NA | 0.005 | 0.954 | -0.004 | 0.979 | -0.145 | 0.800 |
| | G16911 | protoporphyrinogen oxidase | ppox | -0.130 | 0.357 | -0.302 | 0.020 | -0.002 | 0.998 | -1.114 | 0.000 |
| | G17040 | spermine oxidase-like | smo | 0.011 | NA | 0.004 | 0.991 | 0.001 | 0.998 | 0.845 | 0.001 |
| | G17325 | Glycolate oxidase | gox | 0.005 | 0.977 | -0.080 | 0.612 | 0.002 | 0.999 | -1.198 | 0.000 |
| | G17461 | galactose oxidase | gaox | -0.001 | NA | 0.083 | 0.609 | 0.001 | 0.999 | -1.157 | 0.000 |
| | G21130 | sulfite oxidase, mitochondrial | suox | -0.002 | NA | -0.085 | 0.604 | 0.010 | 0.966 | -1.212 | 0.000 |
| | G21313 | putative aldehyde oxidase 2 | aox2 | -0.004 | NA | -0.002 | 0.994 | 0.003 | 0.992 | -1.781 | 0.000 |
| | G21371 | FAD-linked sulfhydryl oxidase | erv1 | -0.002 | NA | 0.039 | 0.528 | 0.005 | 0.822 | 1.368 | 0.033 |
| Photosynthesis | G14978 | ribulose- bisphosphate carboxylase oxygenase large subunit n- isoform 1 | rbcl1 | -0.01274 | 0.83146 | 0.02008 | 0.93665 | 0.00244 | 0.99587 | 0.14008 | 0.63766 |
| | Plast_rbcL | ribulose-1,5-bisphosphate carboxylase/oxygenase large subunit | rbcl2 | 0.13515 | 0.27516 | -0.04067 | 0.84221 | 0.00950 | 0.95970 | -3.66741 | 0.00000 |
| | Plast_rbcS | ribulose-1,5-bisphosphate carboxylase/oxygenase small subunit | rbcs3 | 0.09505 | 0.46121 | -0.02802 | 0.89133 | 0.00365 | 0.99180 | -3.31163 | 0.00000 |
| | Plast_psbE | photosystem II protein V | psII1 | 0.10741 | 0.41939 | -0.02246 | 0.90590 | -0.00742 | 0.98060 | -3.17961 | 0.00000 |
| | Plast_psbF | photosystem II protein VI | psII2 | 0.11463 | 0.33305 | -0.02946 | 0.86955 | -0.00093 | 0.99844 | -3.25821 | 0.00000 |
| Electron chain transport | G1676 | NADH dehydrogenase 1 alpha subcomplex | ndh1 | -0.05415 | 0.65954 | -0.04839 | 0.79978 | -0.00148 | 0.99180 | -1.65974 | 0.00000 |
| | G2076 | NADH dehydrogenase [ubiquinone] 1 alpha subcomplex subunit 6 | ndh2 | -0.00753 | 0.96696 | -0.15422 | 0.31496 | 0.02585 | 0.70949 | 0.39421 | 0.02900 |
| | G2959 | NADH dehydrogenase [ubiquinone] iron-sulfur protein 7, mitochondrial | ndh3 | -0.03181 | 0.82291 | -0.36856 | 0.00525 | 0.01745 | 0.92140 | -1.70821 | 0.00000 |
| | G3901 | NADH dehydrogenase iron-sulfur protein 4 | ndh4 | -0.00043 | 0.99807 | -0.15821 | 0.32674 | -0.01125 | 0.99655 | -0.68036 | 0.00033 |
| | G5588 | NADH dehydrogenase | ndh5 | 0.00326 | NA | -0.00598 | 0.89158 | -0.00044 | NA | 2.11610 | 0.01241 |
| | G8744 | NADH dehydrogenase [ubiquinone] 1 alpha subcomplex subunit 8-B | ndh6 | 0.02283 | 0.88138 | -0.17581 | 0.23662 | 0.01059 | 0.99180 | -0.00888 | 0.96987 |
| | G11240 | NADH dehydrogenase isoform 2 | ndh7 | -0.00392 | 0.98022 | -0.07496 | 0.62344 | 0.02034 | 0.85707 | -2.06155 | 0.00000 |
| | G12211 | NADH dehydrogenase [ubiquinone] 1 alpha subcomplex assembly factor 3 | ndh8 | -0.01961 | NA | -0.00435 | 0.98952 | 0.00312 | 0.99180 | -0.77690 | 0.02552 |
| | G16887 | NADH dehydrogenase i iron-sulfur protein 13kda subunit | ndh9 | -0.00350 | 0.97804 | -0.09544 | 0.52115 | 0.01596 | 0.70297 | -0.62537 | 0.06971 |
| | G18550 | NADH dehydrogenase [ubiquinone] 1 alpha subcomplex subunit 8-B | nndh10 | 0.00649 | NA | -0.09202 | 0.48022 | 0.00284 | 0.99180 | 0.62663 | 0.09656 |

**Table S1.** continuation

| | | | | A+B vs A early exponential | | A+B vs A late exponential | | A+B vs A stationary | | A+B vs A demise | |
|---|---|---|---|---|---|---|---|---|---|---|---|
| Function | Gene ID | Annotation | Short name | logFC | padj | logFC | padj | logFC | padj | logFC | padj |
| Electron chain transport | G22664 | NADH dehydrogenase | ndh11 | 0.00711 | 0.96890 | -0.19067 | 0.09296 | -0.00301 | 0.99417 | -0.84282 | 0.00000 |
| | G26387 | NADH dehydrogenase subunit 1 (mitochondrion) | ndh12 | 0.10895 | 0.40996 | -0.01586 | 0.94227 | 0.01152 | 0.91706 | 1.69196 | 0.00000 |
| | G28238 | NADH dehydrogenase subunit 7 (mitochondrion) | ndh13 | -0.01069 | 0.96040 | -0.06610 | 0.65710 | 0.00393 | 0.99549 | -0.04005 | 0.79348 |
| | G29626 | NADH dehydrogenase iron-sulfur protein 3, mitochodrial | ndh14 | -0.00153 | 0.99665 | -0.08213 | 0.62344 | 0.00595 | 0.98610 | -0.88407 | 0.00000 |
| | G29857 | NADH dehydrogenase [ubiquinone] 1 beta subcomplex subunit 9-like | ndh15 | 0.00358 | 0.99001 | -0.12457 | 0.44189 | -0.00055 | 0.99844 | 0.75985 | 0.00032 |
| | Mito_nad1 | NADH dehydrogenase subunit 1 | ndh16 | 0.17475 | 0.14458 | -0.00658 | 0.97876 | 0.01254 | 0.86951 | 0.58391 | 0.00004 |
| | Mito_nad2 | NADH dehydrogenase subunit 2 | ndh17 | 0.09013 | 0.47017 | -0.02788 | 0.88218 | 0.01441 | 0.83212 | 0.35724 | 0.02986 |
| | Mito_nad3 | NADH dehydrogenase subunit 3 | ndh18 | 0.05248 | 0.66703 | 0.01756 | 0.93083 | 0.00164 | 0.99587 | 1.78672 | 0.00000 |
| | Mito_nad4 | NADH dehydrogenase subunit 4 | ndh19 | 0.15951 | 0.23109 | -0.04063 | 0.76567 | 0.01237 | 0.80355 | 0.39933 | 0.01046 |
| | Mito_nad4L | NADH dehydrogenase subunit 4L | ndh20 | 0.07125 | 0.61769 | 0.01360 | 0.99523 | 0.01294 | 0.87939 | 0.95221 | 0.00000 |
| | Mito_nad5 | NADH dehydrogenase subunit 5 | ndh21 | 0.14681 | 0.25255 | -0.06245 | 0.69821 | -0.00233 | 0.84556 | 1.09112 | 0.00000 |
| | Mito_nad6 | NADH dehydrogenase subunit 6 | ndh22 | 0.10261 | 0.44689 | -0.03348 | 0.84681 | 0.01462 | 0.88280 | 0.45214 | 0.02928 |
| $H_2O_2$ degradation | G14904 | putative peroxidase/catalase | kat1 | -0.007 | NA | -0.024 | 0.918 | 0.011 | 0.938 | -0.277 | 0.138 |
| | G26438 | putative peroxidase/catalase | kat2 | 0.073 | 0.386 | -0.102 | 0.501 | 0.016 | 0.907 | -0.301 | 0.318 |
| | G16849 | ascorbate peroxidase | apx | 0.025 | 0.820 | 0.022 | 0.921 | 0.010 | 0.921 | -0.458 | 0.017 |
| | G19475 | cytochrome C peroxidase | ccp1 | 0.019 | 0.885 | -0.026 | 0.916 | 0.004 | 0.999 | -0.666 | 0.001 |
| | G11059 | cytochrome C peroxidase | ccp2 | 0.033 | NA | -0.087 | 0.069 | 0.000 | NA | 4.774 | 0.000 |
| | G11259 | cytochrome C peroxidase | ccp3 | -0.001 | NA | -0.057 | 0.725 | -0.007 | 0.977 | -1.156 | 0.000 |
| | G9921 | cytochrome C peroxidase | ccp4 | 0.010 | NA | 0.104 | 0.501 | 0.009 | 0.875 | -0.061 | 0.884 |
| | G16606 | putative cytochrome c peroxidase | ccp5 | 0.006 | NA | -0.009 | 0.969 | -0.011 | 0.938 | 0.976 | 0.000 |
| | G26763 | putative cytochrome c peroxidase | ccp6 | 0.020 | 0.834 | -0.042 | 0.817 | -0.003 | 0.998 | 1.006 | 0.000 |
| | G1678 | glutathione peroxidase | gpx1 | -0.006 | 0.920 | -0.148 | 0.364 | 0.009 | 0.921 | -0.737 | 0.000 |
| | G17236 | glutathione peroxidase | gpx2 | -0.009 | 0.906 | -0.114 | 0.444 | -0.016 | 0.729 | -2.153 | 0.000 |
| | G29707 | glutathione peroxidase | gpx3 | -0.003 | 0.987 | -0.104 | 0.514 | 0.002 | 0.995 | -2.730 | 0.000 |

**Table S2. Bacterial genes involved in $H_2O_2$ metabolism.** Log2 fold changes (logFC) in gene expression for *P. inhibens* grown in pure cultures (B) or in co-culture with algae (B+A) across different growth phases. Exp: exponential phase, stat: stationary phase. padj refers to the adjusted *p*-value calculated using the Bonferroni correction. The underlying dataset was previously described in Sperfeld & Narváez-Barragán *et al*.[1]

| | | | | B+A_early exp vs B exp | | B+A late exp vs B exp | | B+A early stat vs B stat | | B+A late stat vs B stat | |
|---|---|---|---|---|---|---|---|---|---|---|---|
| **Function** | **GeneID** | **Annotation** | **Short name** | **logFC** | **padj** | **logFC** | **padj** | **logFC** | **padj** | **logFC** | **padj** |
| $H_2O_2$ tolerance | PGA1_RS19315 | hydrogen peroxide-inducible genes activator | OxyR | -0.764 | 0.466 | -2.627 | 0.002 | -0.464 | 0.704 | 0.148 | 0.999 |
| | PGA1_RS19320 | catalase/peroxidase HPI | KatG | -0.183 | 0.027 | -0.023 | 0.807 | 0.049 | 0.459 | 0.248 | 0.001 |
| | PGA1_RS07285 | peroxiredoxin | Pxn1 | 0.430 | 0.012 | 0.964 | 0.000 | 1.465 | 0.043 | 0.268 | 0.999 |
| | PGA1_RS16395 | peroxiredoxin | Pxn2 | -0.030 | 0.598 | -0.223 | 0.026 | 0.302 | 0.000 | 0.076 | 0.564 |
| | PGA1_RS14545 | peroxiredoxin | Pxn3 | 0.160 | 0.057 | 0.378 | 0.000 | 0.002 | 0.978 | -0.073 | 0.683 |
| | PGA1_RS08660 | peroxiredoxin | Pxn4 | 0.106 | 0.219 | -0.117 | 0.649 | -0.243 | 0.029 | -0.332 | 0.002 |
| | PGA1_RS06550 | cytochrome-c peroxidase | CCP | 0.869 | 0.041 | 0.879 | 0.039 | 0.963 | 0.066 | 0.325 | 0.999 |
| | PGA1_RS10565 | peroxidase-related enzyme | Px-r1 | -0.106 | 0.738 | 0.069 | 0.807 | 0.155 | 0.217 | -0.052 | 0.999 |
| | PGA1_RS14495 | peroxidase-related enzyme | Px-r2 | -0.063 | 0.683 | 0.125 | 0.389 | 0.027 | 0.856 | 0.007 | 0.999 |
| | PGA1_RS15845 | peroxide stress protein YaaA | YaaA | -0.067 | 0.828 | -2.331 | 0.001 | -0.665 | 0.312 | -0.260 | 0.171 |
| $H_2O_2$ production and betaine degradation | PGA1_RS09500 | sarcosine oxidase subunit gamma | SoxG1 | 0.817 | 0.101 | 1.303 | 0.006 | 1.150 | 0.019 | 0.205 | 0.999 |
| | PGA1_RS09505 | sarcosine oxidase subunit alpha family protein | SoxA1 | 0.952 | 0.019 | 1.393 | 0.001 | 1.239 | 0.005 | 0.073 | 0.999 |
| | PGA1_RS09510 | sarcosine oxidase subunit delta | SoxD1 | -0.200 | 0.863 | 1.308 | 0.022 | 1.438 | 0.045 | 0.078 | 0.999 |
| | PGA1_RS09515 | sarcosine oxidase subunit beta family protein | SoxB1 | 1.218 | 0.088 | 1.596 | 0.018 | 1.603 | 0.047 | 0.295 | 0.999 |
| | PGA1_RS18860 | sarcosine oxidase subunit gamma | SoxG2 | -0.386 | 0.755 | -0.202 | 0.844 | 0.572 | 0.305 | 0.041 | 0.999 |
| | PGA1_RS18865 | sarcosine oxidase subunit alpha family protein | SoxA2 | 1.010 | 0.019 | 1.656 | 0.000 | 1.022 | 0.005 | -0.294 | 0.999 |
| | PGA1_RS18870 | sarcosine oxidase subunit delta | SoxD2 | 1.980 | 0.007 | 2.272 | 0.002 | 1.281 | 0.071 | -0.413 | 0.999 |
| | PGA1_RS18875 | sarcosine oxidase subunit beta family protein | SoxB2 | 1.008 | 0.053 | 1.290 | 0.009 | 1.007 | 0.025 | -0.590 | 0.890 |
| | PGA1_RS04345 | sarcosine oxidase subunit beta family protein | SoxB3 | 0.404 | 0.188 | -0.829 | 0.179 | -0.341 | 0.564 | -0.056 | 0.999 |
| | PGA1_RS04350 | sarcosine oxidase subunit delta | SoxD3 | -0.961 | 0.197 | -3.108 | 0.000 | -0.278 | 0.530 | 0.085 | 0.999 |
| | PGA1_RS04355 | sarcosine oxidase subunit alpha family protein | SoxA3 | 0.044 | 0.895 | 0.019 | 0.956 | 0.009 | 0.976 | 0.158 | 0.818 |
| | PGA1_RS04360 | sarcosine oxidase subunit gamma | SoxG3 | -1.206 | 0.201 | -0.730 | 0.389 | 0.212 | 0.520 | 0.137 | 0.999 |
| $H_2O_2$ production | PGA1_RS09485 | superoxide dismutase | SOD | -0.345 | 0.000 | -0.513 | 0.000 | 0.153 | 0.002 | -0.086 | 0.192 |
| | PGA1_RS13820 | L-aspartate oxidase | NadB | 0.201 | 0.123 | 0.502 | 0.000 | 0.242 | 0.568 | 0.008 | 0.999 |
| | PGA1_RS08335 | pyridoxamine 5'-phosphate oxidase | pdxH | 0.105 | 0.389 | 0.302 | 0.012 | 0.113 | 0.358 | -0.105 | 0.872 |
| | PGA1_RS17445 | CopD family protein | hemJ | -1.017 | 0.150 | -3.091 | 0.000 | 0.042 | 0.957 | 0.100 | 0.999 |
| | PGA1_RS18820 | Ldh family oxidoreductase | Ldh | 1.072 | 0.298 | 1.620 | 0.081 | 0.780 | 0.185 | -0.173 | 0.999 |
| | PGA1_RS19020 | dimethylsulfoniopropionate demethylase | dmdA | 0.963 | 0.058 | 1.276 | 0.009 | 1.553 | 0.013 | 0.165 | 0.999 |

**Table S3. Correlations between the expression of *katG* and the DMSP degradation genes *dmdA* and *bmt*.** Correlations were calculated as Pearson correlations with *p*-values adjusted using a false discovery rate of 5%. The underlying dataset was previously described in Sperfeld & Narváez-Barragán *et al*.[1]

| | Pearson's correlation | *P*-value |
|---|---|---|
| *katG vs dmdA* | -0.39 | 0.436 |
| *katG vs bmt* | 0.18 | 0.7306 |

**Table S4. Relative gene expression of genes involved in $H_2O_2$ production in *P. inhibens*, induced by betaine.** Gene expression was measured via qRT-PCR in bacteria cultivated with glucose or betaine as a sole carbon source. Log2fold was calculated as expression induced by betaine, divided by the expression value in glucose.

| Gene | log2fold change | padj |
|---|---|---|
| $sox_1$ | 24.5 | 0.0203 |
| *sod* | 2 | 0.0303 |
| *hemJ* | 1 | 0.01 |
| *pdxH* | 0.464 | 0.0369 |
| *nadB* | 0.224 | 0.0303 |
| *ldh* | 0.261 | 0.15 |
| $sox_2$ | 0.0895 | 0.002 |

### Table S5. Oligonucleotides used in the study.

| Oligonucleotide | Sequence 5’-3’ | Source |
|---|---|---|
| 1.- *katg* upstream forward primer with homology to pCRTM8/GW/TOPO | GTACAAAAAAGCAGGCTCCGAATTCGCCCTTACGCTCACCATGGAAATCAGG | Current study |
| 2.- *katg* upstream reverse primer with homology to gentamycin cassette | CATTACAGTTTACGAACCGAACAGGCTTATGTCAAGTTGTCTCTCCTGGATCTTGG | Current study |
| 3.- *katg* downstream forward primer with homology to gentamycin cassette | TTTGATATCGACCCAAGTACCGCCACCTAATCGGCGCAGATACCATGCAGTC | Current study |
| 4.- *katg* downstream reverse primer with homology to pCRTM8/GW/TOPO | CTTTGTACAAGAAAGCTGGGTCGAATTCGCCCTCCGGGTCCATATCGGCGATG | Current study |
| 5.- *katg* verification forward primer | GACCTCACCGATTTTGCGG | Current study |
| 6.- *katg* verification reverse primer | CTATAGCCTTCGGCACAGAC | Current study |
| 7.- *pxn* upstream forward primer with homology to pCRTM8/GW/TOPO | GTACAAAAAAGCAGGCTCCGAATTCGCCCTTAGCGAATGGAAATAGCGCGCAC | Current study |
| 8.- *pxn* upstream reverse primer with homology to gentamycin cassette | GCATTACAGTTTACGAACCGAACAGGCTTATGTCAACATCGGCTCCTTAAGGATCAATTCG | Current study |
| 9.- *pxn* downstream forward primer with homology to gentamycin cassette | GCACTTTGATATCGACCCAAGTACCGCCACCTAATCCAGAGGGTCTGAGATACCGAC | Current study |
| 10.- *pxn* downstream reverse primer with homology to pCRTM8/GW/TOPO | CTTTGTACAAGAAAGCTGGGTCGAATTCGCCCTGTCACTATGCCGAAGACATGCC | Current study |
| 11.- *pxn* verification forward primer | GGAGATACTGACCGATGGTCAGG | Current study |
| 12.- *pxn* verification reverse primer | GGAGGCGTGCCGATATCTTG | Current study |
| 13.- Gentamycin cassette forward primer | GTTGACATAAGCCTGTTCG | Sperfeld & Narváez-Barragán *et al.*[1] |
| 14.- Gentamycin cassette reverse primer | GTTAGGTGGCGGTACTTGG | Sperfeld & Narváez-Barragán *et al.*[1] |
| 15.- Gentamycin cassette verification forward primer | GTGCAAGCAGATTACGGTGACG | Sperfeld & Narváez-Barragán *et al.*[1] |
| 16.- Gentamycin cassette verification reverse primer | GAGCCTACATGTGCGAATGATGC | Sperfeld & Narváez-Barragán *et al.*[1] |
| 17.- *sod* upstream forward primer with homology to pCRTM8/GW/TOPO | GTACAAAAAAGCAGGCTCCGAATTCGCCCTTACACGATGAATGGACTGCGGAAG | Current study |
| 18.- *sod* upstream reverse primer with homology to kanamycin cassette | GCATTACAGTTTACGAACCGAACAGGCTTATGTCAACTGGGCCCCCTGTAGTATCTG | Current study |
| 19.- *sod* downstream forward primer with homology to kanamycin cassette | GCCTTCTATCGCCTTCTTGACGAGTTCTTCTGATTGCGGCGCAGCAGCATTAG | Current study |
| 20.- *sod* downstream reverse primer with homology to pCRTM8/GW/TOPO | CTTTGTACAAGAAAGCTGGGTCGAATTCGCCCTCTGAAGGCTCTGGTGGAGCATG | Current study |
| 21.- *sod* verification forward primer | GGTGCTGACCTTTTGCTGC | Current study |
| 22.- *sod* verification reverse primer | GGTGATGCTCTACAATGCCG | Current study |
| 23.- Kanamycin cassette forward primer | TTGACATAAGCCTGTTCGGTTCG | Current study |
| 24.- Kanamycin cassette reverse primer | GAGCACGTACTCGGATGGAA | Current study |
| 25.- Kanamycin cassette verification forward primer | GAGCACGTACTCGGATGGAA | Narváez-Barragán *et al.*[3] |
| 26.- Kanamycin cassette verification reverse primer | TTCCATCCGAGTACGTGCTC | Narváez-Barragán *et al.*[3] |
| 27.- $sox_1$ forward primer for qRT-PCR | GGGTGTGCATTTTGAACGCT | Current study |
| 28.- $sox_1$ reverse primer for qRT-PCR | TTTCAGATCCCGCCATGTCC | Current study |
| 29.- *sod* forward primer for qRT-PCR | GACGGCAAGTCCATGGAAGA | Current study |
| 30.- *sod* reverse primer for qRT-PCR | AAGGTGTCAACGGAGCCAAA | Current study |
| 31.- *hemJ* forward primer for qRT-PCR | CGGACAGCCTTCCCATCTTT | Current study |
| 32.- *hemJ* reverse primer for qRT-PCR | ACCAGCAACCAGTGGTGAAA | Current study |
| 33.- *pdxH* forward primer for qRT-PCR | TGGAAATCACTGCGTCGTCA | Current study |
| 34.- *pdxH* reverse primer for qRT-PCR | TTTGACCCCTTGGTTGCAGT | Current study |
| 35.- *nadB* forward primer for qRT-PCR | GCCAGAAGTCTCGCTGAGTT | Current study |
| 36.- *nadB* reverse primer for qRT-PCR | TGGCGCACATATTGGCAAAG | Current study |
| 37.- *ldh* forward primer for qRT-PCR | GACTGCGACAGAAACCCTGA | Current study |
| 38.- *ldh* reverse primer for qRT-PCR | CATCAACCTTGCCGCATTCC | Current study |
| 39.- $sox_2$ forward primer for qRT-PCR | GCCACGGTTGTTGAGCTTTT | Current study |
| 40.- $sox_2$ reverse primer for qRT-PCR | GCGAATGATCGGCTCGTAGA | Current study |
| 41.- *gyrA* forward primer for qRT-PCR | GCCGATTCCTGACCTCCTTC | Lipsman *et al.*[4] |
| 42.- *gyrA* forward primer for qRT-PCR | TCAGCTTATGTCGGGCTTCG | Lipsman *et al.*[4] |
| 43.- *recA* forward primer for qRT-PCR | GCTGACACCCAAGTCGGAG | Lipsman *et al.*[4] |
| 44.- *recA* forward primer for qRT-PCR | AGCCGAACATAACGCCAATCT | Lipsman *et al.*[4] |

**Table S6. List of Multiple reaction monitoring (MRM).** MRMs transitions used for the detection and quantification of betaine by LC-MS/MS.

| Name | Ionization mode | Cone (V) | MRM | CE (eV) | Used for |
|---|---|---|---|---|---|
| Betaine | + | 20 | 118.0 > 58.9 | 18 | Quantification |
| | | | 118.0 > 58.0 | 18 | Identification |
| | | | 118.0 > 42.0 | 25 | |
| | | | 118.0 > 74.0 | 15 | |
| | | | 118.0 > 102.0 | 15 | |
| $^{13}C_5{}^{15}N$-Betaine | + | 20 | 124.0 > 62.0 | 18 | Quantification |
| | | | 124.0 > 63.0 | 18 | Identification |

**References for Supplementary Materials**

1.- Sperfeld, M., Narváez-Barragán, D.A., Malitsky, S., Frydman, V., Yuda, L., Rocha, J., and Segev, E. (2024). Algal methylated compounds shorten the lag phase of Phaeobacter inhibens bacteria. Nat Microbiol, 1–16. https://doi.org/10.1038/s41564-024-01742-6.

2.- Salazar, G., Paoli, L., Alberti, A., Huerta-Cepas, J., Ruscheweyh, H.-J., Cuenca, M., Field, C.M., Coelho, L.P., Cruaud, C., Engelen, S., et al. (2019). Gene Expression Changes and Community Turnover Differentially Shape the Global Ocean Metatranscriptome. Cell *179*, 1068-1083.e21. https://doi.org/10.1016/j.cell.2019.10.014.

3.- Narváez-Barragán, D.A., Sperfeld, M., and Segev, E. DmdA-independent lag phase shortening in Phaeobacter inhibens bacteria under stress conditions. The FEBS Journal. https://doi.org/10.1111/febs.70128.

4.- Lipsman, V., Shlakhter, O., Rocha, J., and Segev, E. (2024). Bacteria contribute exopolysaccharides to an algal-bacterial joint extracellular matrix. npj Biofilms Microbiomes 10, 1–16. https://doi.org/10.1038/s41522-024-00510-y.
